# Arf family GTPases are present in Asgard archaea

**DOI:** 10.1101/2024.02.28.582541

**Authors:** Romana Vargová, Roxanne Chevreau, Marine Alves, Camille Courbin, Kara Terry, Pierre Legrand, Marek Eliáš, Julie Ménétrey, Joel B. Dacks, Catherine L. Jackson

**Affiliations:** Department of Biology and Ecology, Faculty of Science, University of Ostrava, Ostrava, Czech Republic; Division of Infectious Diseases, Department of Medicine, and Department of Biological Sciences, University of Alberta, Edmonton, AB, Canada; Université Paris-Saclay, CEA, CNRS, Institute for Integrative Biology of the Cell (I2BC), 91198, Gif-sur-Yvette, France; Université Paris Cité, CNRS, Institut Jacques Monod, 75013 Paris, France; Synchrotron SOLEIL, l’Orme des Merisiers, 91190, Saint Aubin, France; Institute of Parasitology, Biology Centre, Czech Academy of Sciences, České Budějovice, Czech Republic; Centre for Life’s Origins and Evolution, Department of Genetics, Evolution, & Environment, University College, London

**Author notes:** Co-corresponding authors (Marek Elias,; Julie Ménétrey,; Joel B. Dacks,; Catherine L. Jackson,).

**Keywords:** Eukaryogenesis, GTPase, Arf, Asgard, endoplasmic reticulum, plasma membrane

## Abstract

The emergence of eukaryotes from their prokaryotic ancestors is one of the most fundamental evolutionary events in the history of life. Little is robustly known about how eukaryogenesis occurred, but a major breakthrough came with the identification of the Asgardarchaeota, the closest prokaryotic lineage to eukaryotes yet discovered. Endomembrane organelles, and the capacity to transport material between them, are major hallmarks of eukaryotic cells. The Arf family GTPases are crucial regulators of organelle dynamics in eukaryotes, functioning in vesicle budding, membrane tethering and membrane-cytoskeleton interactions. Although an expanded GTPase complement has been reported in the Asgardarchaeota, the specific origins of the Arf family remain elusive. Here we report a new group of prokaryotic GTPases, the ArfRs. Widely present in Asgardarchaeota and almost exclusive to them, it is the clade from which all eukaryotic Arf family proteins are derived. Heterologous expression of representative Asgardarchaeota ArfR proteins in the model eukaryote *Saccharomyces cerevisiae* and X-ray crystallographic studies demonstrate that ArfR GTPases possess the mechanism of membrane binding and structural features unique to Arf family proteins. Our results show that Arf family GTPases are present in Asgardarchaeota, and strongly suggest that they originated in the archaeal contributor to eukaryogenesis, providing support for nascent endomembrane system capacity evolving early in eukaryogenesis.

## Main

Eukaryogenesis was one of the most transformative evolutionary transitions in the history of life on Earth, setting the stage for nearly all complex life that we see today. The processes and timelines involved, although hotly debated, are poorly understood. It is generally agreed that eukaryotes originated from an archaeal lineage that entered into symbiosis with at least one bacterial lineage. Whereas an alphaproteobacterial contributor, giving rise to the mitochondria, was an essential component in eukaryogenesis, the archaeal host lineage clearly contributed many of the building blocks for the other organelles, including those of the endomembrane system (eg. endoplasmic reticulum, Golgi, endosomes and lysosomes) and the nucleus. Beyond this basic agreement, however, there are numerous contentiously competing hypotheses proposing different processes, bases for interaction, and order in which diagnostic eukaryotic traits arose^1–6^.

Firmly establishing what cellular features originated in the archaeal vs bacterial contributors is one crucial way forward to addressing the critical questions of the origins and relative timing of emergence of the characteristic eukaryotic systems during eukaryogenesis. The discovery of the Asgardarchaeota was revolutionary, providing a target as to the identity of the archaeal lineage contributing to eukaryogenesis^7,8^. While contention does exist, the consensus places eukaryotes within the Asgardarchaeota, with the most recent data concluding the heimdallarchaeial ancestry of eukaryotes^9^. Moreover, the increasing number of genes from Asgardarchaeota that encode proteins previously only seen in eukaryotes, i.e. Eukaryotic Signature Genes (ESGs), provides a separate line of evidence for the Asgardarchaeota as the archaeal contributors to eukaryogenesis^7–10^. The asgardarchaeote ESGs include those encoding proteins of ubiquitylation systems, glycosylation systems (OST complex), various aspects of information processing, actin and tubulin cytoskeleton systems, and multiple membrane trafficking components^7–10^. Notably, the distribution of the encoded components is not homogenous across the Asgards, implying some combination of horizontal gene transfer (HGT) and gene loss shaping the repertoire^9,10^ and thus mitigating emphasis on any one particular asgardarchaeote lineage. Nonetheless, just as mitochondrial traits found in the Alphaproteobacteria suggest their origins from this bacterial contributor, aspects of eukaryotic cell biology found within the Asgardarchaeota provide specific evidence for their archaeal origin, and therefore likely present prior to endosymbiosis in at least some nascent form.

Another point that is largely agreed on amongst eukaryogenesis theories is the essentiality of evolving a system for cellular trafficking, i.e. a system that allows compartmentalization and targeted intracellular movement, facilitating cellular size increase. Not only must the emergence of components of such a system be explained in any model of evolutionary transition, the implementation of such a system is a critical and common aspect even of competing mathematical models of energetic considerations in eukaryogenesis^11–13^. Such a system in eukaryotes involves both the cytoskeleton and membrane-trafficking machinery, homologues of which have been identified in Asgardarchaeota. These include SNARE domains, ESCRT complex I, II and III components, TRAPP complex homologues, COPII vesicle coat components, multiple domains of adaptin coat subunits, longin domains, DENN domains, and an expanded repertoire of small G proteins^7–10,14,15^.

Although evolutionary homology demonstrates origins, and can be indicative of function, assuming conservation, there are limits to what bioinformatics can predict. On the timescale and likely morphological and ecological differences between Asgardarchaeota and eukaryotes, the functional homology of ESGs is far less certain than their ancestry. Elegant work addressing this gap has been obtained through structural and biochemical characterization of asgardarchaeote actin and several of its regulators^16–19^, as well as tubulin, which was shown to form microtubule-like filaments^20^. These results strongly argue for the presence of a dynamic cytoskeleton in the archaeal contributor to eukaryogenesis, allowing for cell shape modulation and cytoplasmic trafficking. However, cytoskeleton alone is insufficient for membrane trafficking or an endomembrane system.

In eukaryotes, numerous proteins assure the functioning of the endomembrane system, and in particular membrane trafficking. The best-studied mechanism of membrane trafficking is the formation of transport vesicles from a donor compartment and their targeting and fusion to an acceptor compartment. There are two families of GTPases in eukaryotes whose major functions are in membrane remodelling and trafficking, the Arf and Rab families. Phylogenetic analyses show that Rabs are more closely related to the Ras, Rho, or Ran family than to the Arf family and share with them a separate prokaryotic, most likely asgardarchaeote, ancestor^21,22^. The origin of the Arf family is less well understood, but it was already well-established in the last eukaryotic common ancestor (LECA), with 15 members present^23^. Arf family proteins, including classical Arfs as well as a plethora of Arf-like (Arl) proteins, Arfrp1 and Sar1, function in multiple steps of membrane trafficking including vesicle budding, movement of vesicles and organelles along cytoskeleton tracks, targeting and regulation of vesicle fusion, as well as in membrane lipid modification, lipid metabolism, cytoskeleton-membrane interaction and cilia transport^24–26^. Like other membrane-associated small GTPases in eukaryotes, Arf family proteins carry out their functions by recruiting effectors to the membrane in their active GTP-bound form^24,25,27^.

The defining feature of Arf family proteins, which distinguishes them from other GTPases, is the presence of an N-terminal amphipathic helix (AH) that changes conformation upon nucleotide exchange^28–31^. Like other small G proteins, Arf family proteins change conformation in their switch regions upon release of GDP and binding of GTP, but in addition, they undergo a second conformational change by which the N-terminal AH is released from a hydrophobic pocket and associates with membranes. This mechanism was deduced by biochemical^28^ and structural studies, the latter demonstrating that the hydrophobic pocket into which the N-terminal AH is bound in the GDP-bound form^29,32,33^ is occluded in the GTP-bound form by the movement of the interswitch region upon release of GDP^34^. The crystal structures of numerous eukaryotic Arf family proteins have been solved, and all of them exhibit this characteristic conformational change between GDP– and GTP-bound forms^34–39^.

Essentially all membrane trafficking routes within eukaryotic cells are regulated by at least one Arf family protein, and the wide range of functions of these GTPases allows vesicular trafficking to be coordinated with diverse cellular processes^25,27^. The origins of the Arf family GTPases, and the time of emergence of their hallmark mechanism of membrane localization, are currently unknown. To address this gap in understanding, we searched for specific relatives of the Arf GTPases amongst prokaryotes.

### Arf family proteins in Asgardarchaeota

Our previous interest in the phylogeny and diversity of the Arf family in eukaryotes^23^, led us to identify a protein (Kcr_0908; GenBank ACB07654.1) encoded by the genome of *Candidatus* Korarchaeum cryptofilum, a member of the class Korarchaeia in archaeal phylum Thermoproteota (also known as the TACK superphylum)^40^, as more similar to eukaryotic Arf family members than any prokaryotic homolog in the databases at that time. An initial phylogenetic analysis, incorporating Ras superfamily GTPases from the first reported asgardarchaeote metagenome-assembled genomes (MAGs)^7,8,41^, indicated that this korarchaeial GTPase is part of a strongly supported clade within the Ras superfamily tree together with a few unclassified metagenomic sequences, various GTPases from two asgardarchaeote subgroups now called Lokiarchaeales and Heimdallarchaeia, and eukaryotic Arf family members plus SRβ (the eukaryote-specific beta subunit of the signal recognition particle receptor; Extended Data Fig. 1). Crucially, these Arf-related archaeal GTPases were distinct from the previously defined prokaryotic Ras superfamily subgroups, including MglA, Rup1, Rup2, RasL I-IV, Rag, and ArfL^22^. We hypothesized that these archaeal GTPases represent the closest relatives of the eukaryotic Arf/SRβ proteins and constitute a novel subgroup that we denoted ArfR (for “Arf-related”). Notably, the korarchaeial ArfR protein was noticed before but its relationship to the eukaryotic Arf family proteins was missed^42^.

To more systematically assess ArfR distribution and its evolutionary relationship to eukaryotic GTPases, we searched for the occurrence of proteins with a putative specific relationship to the eukaryotic Arf/SRβ proteins in Archaea. Notably, putative ArfR proteins outside Asgardarchaeota were only identified in Korarchaeia; other candidates turned out to represent taxonomically misidentified asgardarchaeote or eukaryote sequences (Table S1). Hence, we focused further analyses on genome assemblies from Asgardarchaeota. Given the nature of the data concerned (primarily MAGs of varying quality, rapidly accumulating in databases and with inconsistent classification across studies), we performed our own survey of the available genome data and selected 62 Asgard genome assemblies making a rich and phylogenetically balanced sample of the asgardarchaeote phylogenetic diversity as recorded in public repositories at the time of our analyses (Fig. S1; Table S2). In fact, our approach enabled us to sample representatives of all known major asgardarchaeote lineages, including a novel class-level lineage that was recognized only recently and denoted Asgardarchaeia^9^. We then searched these genomes with BLAST using as queries the sequences of the human Arf1 and SRβ, along with representatives of ArfR and the previously defined major groups of asgardarchaeote Ras superfamily GTPases (ArfL, RasL, RagGtr, MglA1 and MglA2-5). Significant hits were parsed and manually curated to generate high-quality sets of Ras superfamily sequences for each asgardarchaeote genome, altogether amounting to 2922 sequences (Table S3).

Sequence similarity-based clustering was used to get a picture of the diversity of asgardarchaeote Ras superfamily proteins and their relationship to eukaryotic homologs. Highlighting the previously identified representatives of the different major GTPase subgroups revealed they are separated according to these subgroups, and mark different broader clusters also containing additional (i.e., newly analysed) sequences (Extended Data Fig. 2). Consistent with phylogenetic analyses, the asgardarchaeote RasL sequences were contained with no evident separation into one prominent cluster together with eubacterial Rup1 sequences and Rab/Ras/Rho/Ran GTPases from eukaryotes. The asgardarchaeote and eukaryotic RagGtr proteins formed a less compact cluster, reflecting a higher sequence divergence between different members of this family. Crucially, ArfR sequences belonged to a cluster clearly separate from those corresponding to the previously defined prokaryotic small GTPase groups (including ArfL), and members of the eukaryotic Arf family plus SRβ clustered together with ArfR sequences – closely to them or as part of looser extensions of this cluster, the latter being formed by the more divergent subfamilies Arl16 and SRβ. A number of individual asgardarchaeote small GTPase sequences or small clusters were more or less distantly separated from the annotated clusters and represent divergent offshoots of the major GTPase groups or taxon-specific novel groups akin to the previously defined Rup2 group (which is restricted to a single family of the asgardarchaeote sister group Thermoproteota). One of these, here denoted Rup3, evolutionarily separated from other GTPase groups and characterized by unusual expansions within the GTPase domain (Extended Data Fig. 1 and 2; Fig. S2), was particularly prominent and represented by genes from across the class Heimdallarchaeia. No potential eukaryotic relatives of Rup3 were identified, pointing to a unique feature of the heimdallarchaeial cell biology that was not passed to the descending eukaryotic cell.

The recognition of ArfR as a distinct group of prokaryotic small GTPases was supported by phylogenetic analysis of a large dataset of archaeal sequences (Fig. S3). To allow for more sophisticated phylogenetic analyses needed to address the specific relationship of archaeal and eukaryotic Ras superfamily members, we applied the ScrollSaw procedure^43^ to further reduce the size of the archaeal small GTPase dataset by selecting the most slowly evolving representative sequences (see Materials and Methods for further explanation). The phylogenetic tree inferred from a combination of the reduced archaeal dataset with eubacterial and eukaryotic small GTPase representatives provided robust support for the delineation of the major Ras superfamily lineages (Fig. 1; Fig. S4). Thus, apart from the exclusively prokaryotic MglA1 and MglA2-5 groups and the exclusively asgardarchaeote ArfL group, three additional clades were resolved, each combining sequences from both prokaryotes and eukaryotes. Rab/Ras/Rho/Ran GTPases (as well as the eubacterial Rup1 and Thermoproteaceae-specific Rup2 groups) were embedded within a clade together with asgardarchaeote RasL sequences. Notably, we did not recover any asgardarchaeote sequences specifically sister to eukaryotic Rab GTPases, as the latter formed a clade together with Ras and Rho family sequences; this is counter to reports of *de facto* Rabs being present in Asgardarchaeota^44^. The RagGtr clade contained eukaryotic sequences (two paralogous lineages preceding the LECA) nested among asgardarchaeote relatives. Finally, the eukaryotic Arf family and SRβ sequences were part of a highly supported clade with asgardarchaeote and two representative korarchaeial sequences annotated by us as ArfR. Notably, SRβ was nested among Arf family members in this analysis, consistent with previous structural evidence indicating their specific relationships^45^ and speaking for inclusion of SRβ into the Arf family.

**Figure 1.**
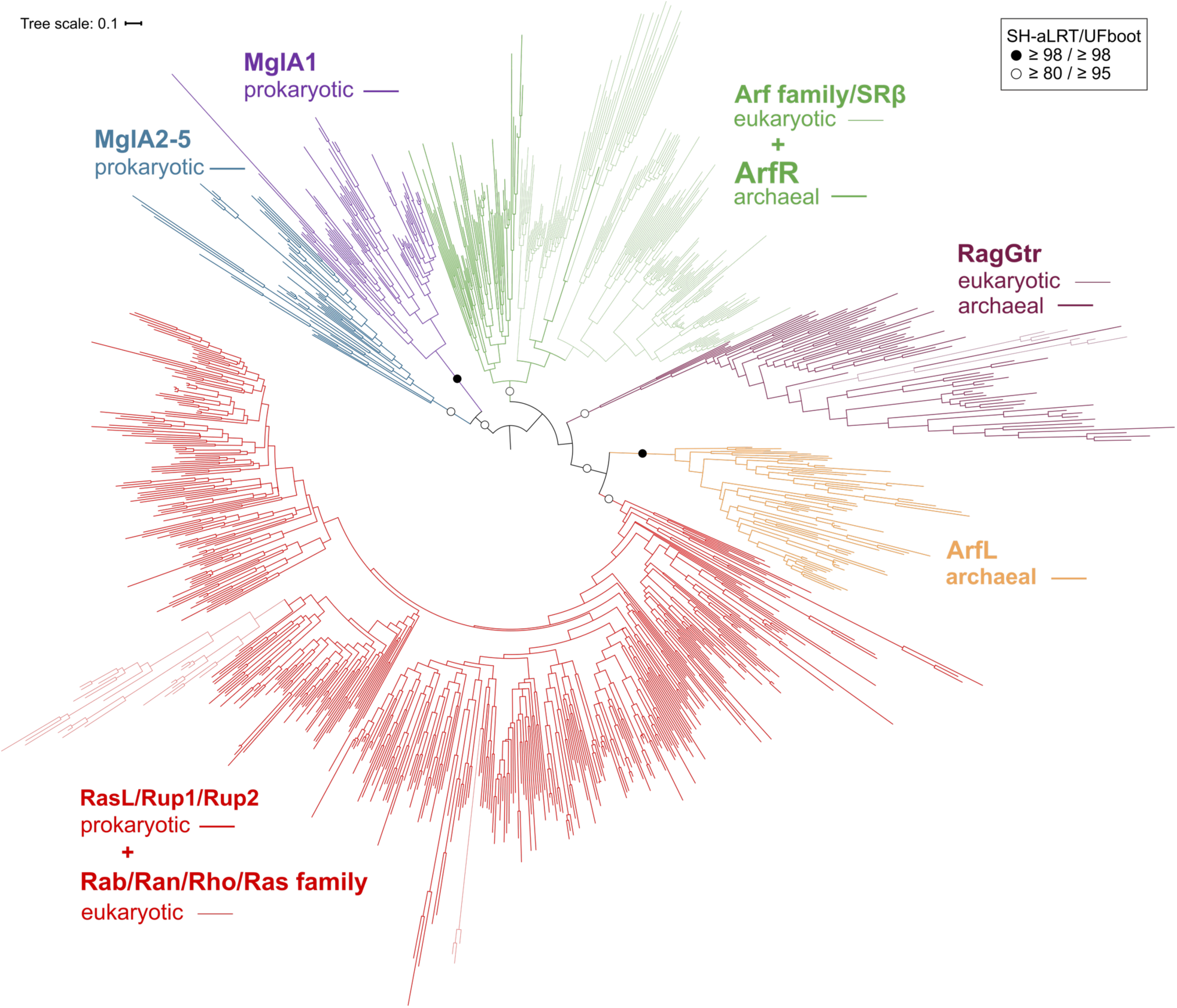
Phylogenetic analysis of a representative dataset of the Ras superfamily sequences from Archaea and reference eukaryotic and eubacterial sequences. The tree was inferred using IQ-TREE with the LG+I+G4 model from a multiple alignment of 996 sequences. Reference eukaryotic and prokaryotic Ras superfamily sequences were selected based on those used in Klinger et al. (2016) and Vargová et al. (2021)^22,23^. Statistical support as assessed with the SH-aLRT test and the ultrafast bootstrap algorithm (both 1000 replicates) is indicated for the deepest branches. The root of the tree was placed between MglA sequences (distributed in two deeply separated subgroups, MglA1 and MglA2-5; see also Klinger et al. 2016^22^), following the results of a broader analysis including EF-Tu as an outgroup (Extended Data Fig. 1). The figure displays a simplified version of the tree (its full version, including tip labels and support values for all internal branches, is provided as Fig. S4).

A closer look at the ArfR occurrence in Asgardarchaeota revealed a patchy distribution previously noticed to be common for many other ESGs and exhibited also by ArfL (Fig. 2). Specifically, with the current sampling and data available, ArfR genes seem to be either completely missing from the given major asgardarchaeote taxon or are common in the taxon representatives. However, considering that according to the most recent inference the immediate archaeal ancestor of eukaryotes belonged to Heimdallarchaeia^9^, it is notable that ArfR is nearly ubiquitous (being absent only from most Njordarchaeales members) in the heimdallarchaeial representatives sampled. To shed more light on the origin of the korarchaeial ArfR genes, we analysed all available korarchaeial genome assemblies and found that ArfR is restricted to a terminal branch shallowly nested in the korarchaeial radiation (Fig. S5). All korarchaeial ArfR genes form a clade nested among asgardarchaeote homologs (Fig. S6), implying that a relatively recent horizontal transfer of a single ArfR gene from an asgardarchaeote donor into a specific korarchaeial lineage is the more likely explanation of the occurrence of ArfR in Korarchaeia than the emergence of ArfR before the Asgardarchaeota-Thermoproteota split. Interestingly, the very first ArfR gene to be encountered (the one from *Ca.* Korarchaeum cryptophilum) happened to be a rare “second-hand” exemplar.

**Figure 2.**
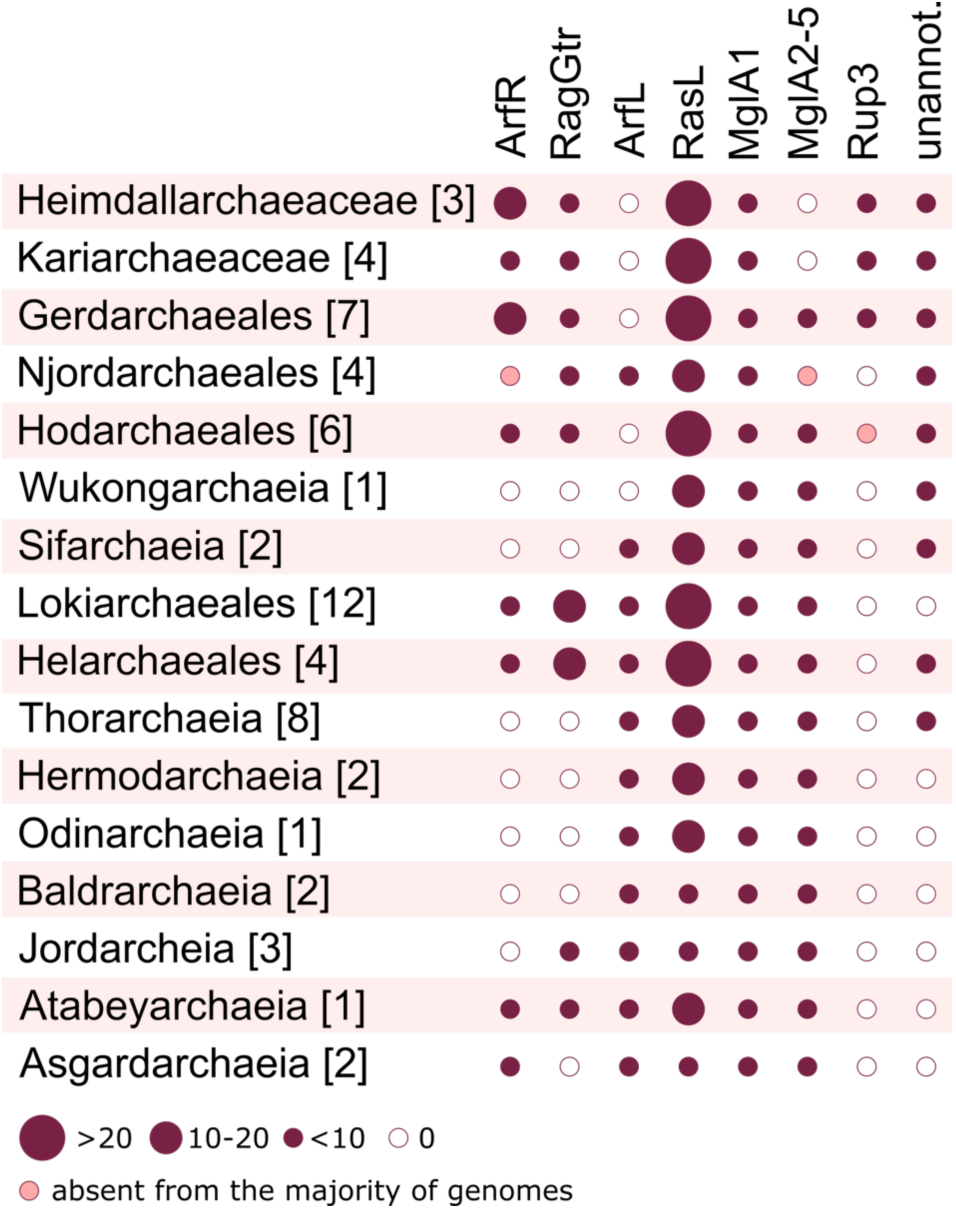
Distribution of the major subgroups of the Ras superfamily across different lineages of Asgardarchaeota. The asgardarchaeote taxa are defined according to Eme et al. (2023)^9^, with the two “Sigynarchaeota” genomes analysed here included in Lokiarchaeales. The numbers in square brackets correspond to the number of genome assemblies selected for analysis for each taxon (listed in Table S2). The circles of a varying size (see graphical legend on the bottom for the scale) shown for each combination of a taxon and a Ras superfamily subgroup represent the median of the number of genes belonging to the subgroup per genome. The small pink circles indicate GTPase subgroups present in only one genome out of multiple analysed here. The last column (labelled “unannot.”) correspond to Ras superfamily genes that were not assigned to any of the separately shown subgroup.

We thus posit that ArfR is an Asgardarchaeota-specific evolutionary innovation that founded the Arf family as a whole and is directly ancestral to eukaryotic Arf family members (including SRβ) via vertical gene inheritance by the eukaryotic lineage from its asgardarchaeote (hemdallarchaeial) progenitor. A previous study did report the presence of “Arf-like” GTPases in Asgardarchaeota, but these either corresponded to the ArfL group^22^ that we show here is in fact not directly related to the eukaryotic Arf family proteins, or the assignment was based only on sequence similarity analyses without investigating the relationship of the asgardarchaeote and eukaryotic proteins by phylogenetic methods^8,10^, which is rectified by our work. Unfortunately, we were unable to robustly resolve whether all eukaryotic members of the Arf family trace their origin to a single ArfR gene inherited by eukaryotes from their asgardarchaeote ancestors or whether different family members evolved from different ArfR ancestors, even despite analyses of a reduced dataset focused on ArfR plus the eukaryotic sequences (data not shown; see also legend to Fig. S4).

### Asgard ArfR localisation in cells

The hallmark of eukaryotic Arf family proteins is the N-terminal AH upstream of the GTPase domain which mediates membrane binding in a nucleotide-dependent manner. To investigate whether the ArfR proteins have this signature property of eukaryotic Arf family members, we searched within the Asgardarchaeota ArfR proteins for those predicted to have an N-terminal AH. Among ninety-four candidates selected for analysis (see Methods), forty-five were identified with a predicted N-terminal AH (Table S3). Among these candidates, two were chosen for further analysis, one from Hodarchaeales (HodArfR1) and one from Gerdarchaeales (GerdArfR1). Neither of these ArfR proteins has the glycine in position 2 that would be required for myristoylation. HodArfR1 and GerdArfR1 share 36% sequence identity, and similarly they share respectively 37% and 39% sequence identity with human Arf1 (for comparison human Arf1 shares 52% sequence identity with the parasitic *Leishmania major* Arl1, while only 30% with human Arl8A).

Because no organisms from Heimdallarchaeia have been cultivated, it is impossible to study ArfR localization in the native cellular context. Therefore, heterologous expression was undertaken in a well-characterized model for studying Arf family function, *Saccharomyces cerevisiae*. To determine whether these proteins have the capacity to bind membranes in cells, synthetic genes encoding the HodArfR1 and GerdArfR1 proteins were inserted into a yeast expression vector containing a GFP coding sequence to allow expression of the Asgard proteins fused to GFP. Confocal microscopy was used to determine the localization of the ArfR-GFP proteins in yeast cells. HodArfR1-GFP clearly localised to yeast membranes, primarily to the endoplasmic reticulum (ER) and the plasma membrane (PM) (Fig. 3a, Extended Data Fig. 3). In yeast, as in all eukaryotes studied to date, the ER and PM have very different membrane compositions^46,47^. These results indicate that the ArfR proteins do not target a specific membrane in yeast cells, but can associate with membranes of different lipid composition. The lack of localization specificity by the asgardarchaeote ArfR proteins suggests that any eukaryotic organelle specificity arose in lineages post-dating the Asgard ancestor of eukaryotes.

**Figure 3.**
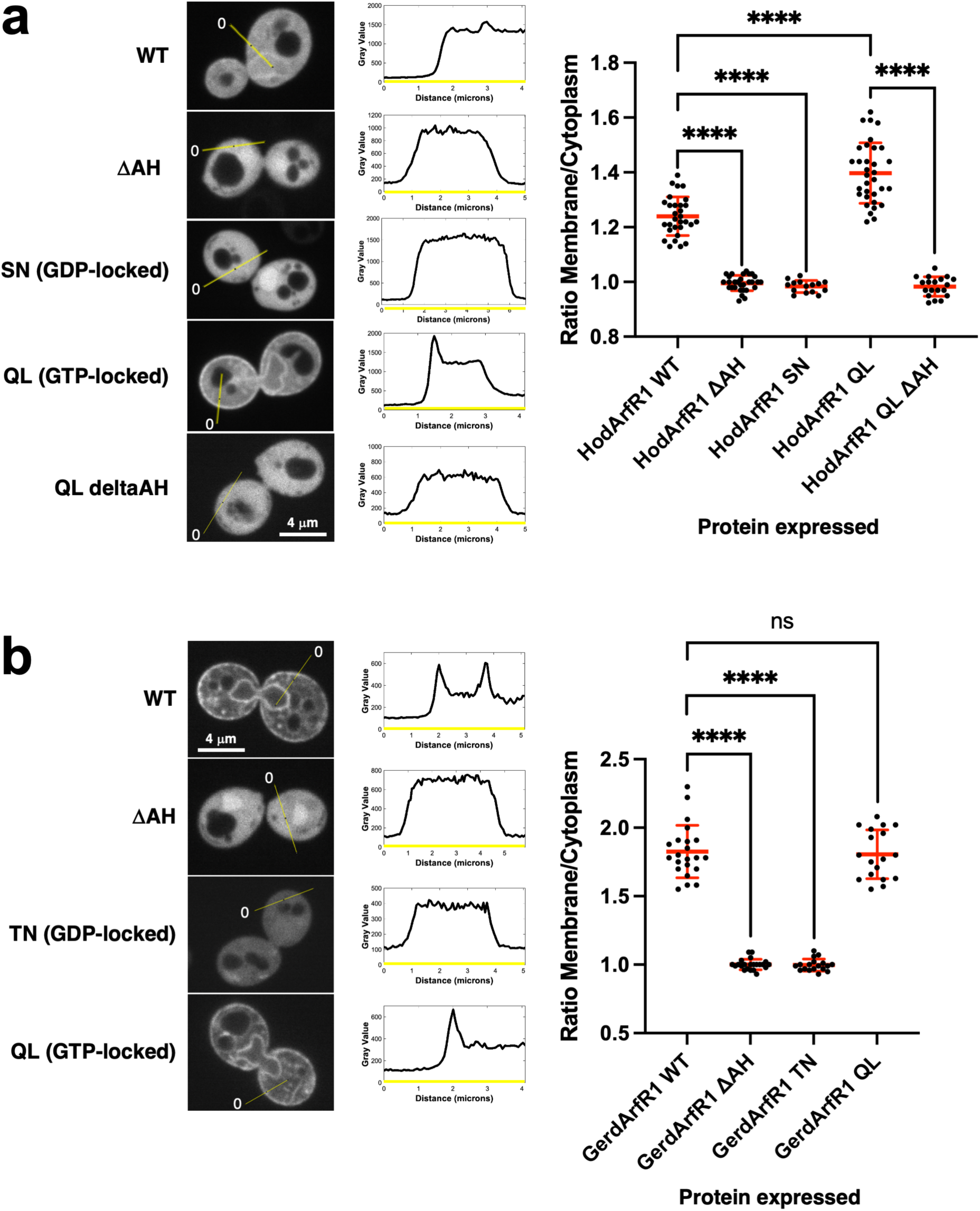
Localization of asgardarchaeote ArfR proteins in yeast cells. HodArfR1 (a) and GerdArfR1 (b) were expressed as GFP fusion proteins in *Saccharomyces cerevisiae* cells. The localizations of wild type and the indicated mutants were quantified as described in Methods. Representative images for each protein expressed in yeast are shown (left panels). The yellow lines are those used to generate plots of pixel intensities (middle panels) to determine membrane/cytosol ratios for each cell analysed (right panels, see Methods for details). “0” (right panels) indicates the beginning of the line shown, and corresponds to “0” on the x-axis of the plots (middle panels). WT, wild type; βAH, mutant with deletion of the N-terminal amphipathic helix; SN or TN, dominant negative mutations HodArfR1-S31N and GerdArfR1-T27N; QL, constitutively active mutations HodArfR1-Q72L and GerdArfR1-Q69L.

We next assayed the localization of an N-terminally truncated form of the HodArfR1-GFP. Importantly, deletion of the N-terminal AH completely abolished membrane binding (Fig. 3a), indicating that the AH is required for association with membranes in cells. Classic GTPase mutations that block GTP binding (S/T31N) or block GTP hydrolysis (Q72L) were introduced into HodArfR1-GFP. The predicted GTP binding-defective S31N mutant was completely cytosolic, whereas the constitutively active HodArfR1-Q72L mutant bound to both ER and plasma membranes (Fig. 4a, Extended Data Fig. 3) and in fact had a significantly higher membrane/cytosol ratio than the wild type (WT) HodArfR1 protein (Fig. 3a). Hence in yeast cells, the HodArfR1 mutant that is unable to bind GTP is cytosolic, whereas the HodArfR1mutant that binds GTP constitutively has a higher level of membrane binding than the WT protein. These results support the conclusion that the GTPase cycle regulates membrane binding of HodArfR1 in the same manner as for eukaryotic Arf family proteins^48^. Because membrane binding of the Q72L mutant was higher than for the WT, we deleted the N-terminal AH in this mutant. Similar to the WT protein with the N-terminal AH deleted, no membrane binding was detected (Fig. 3a, Fig. 4a), strengthening the conclusion that the N-terminal AH is required for binding of the protein to membranes in yeast cells.

**Figure 4.**
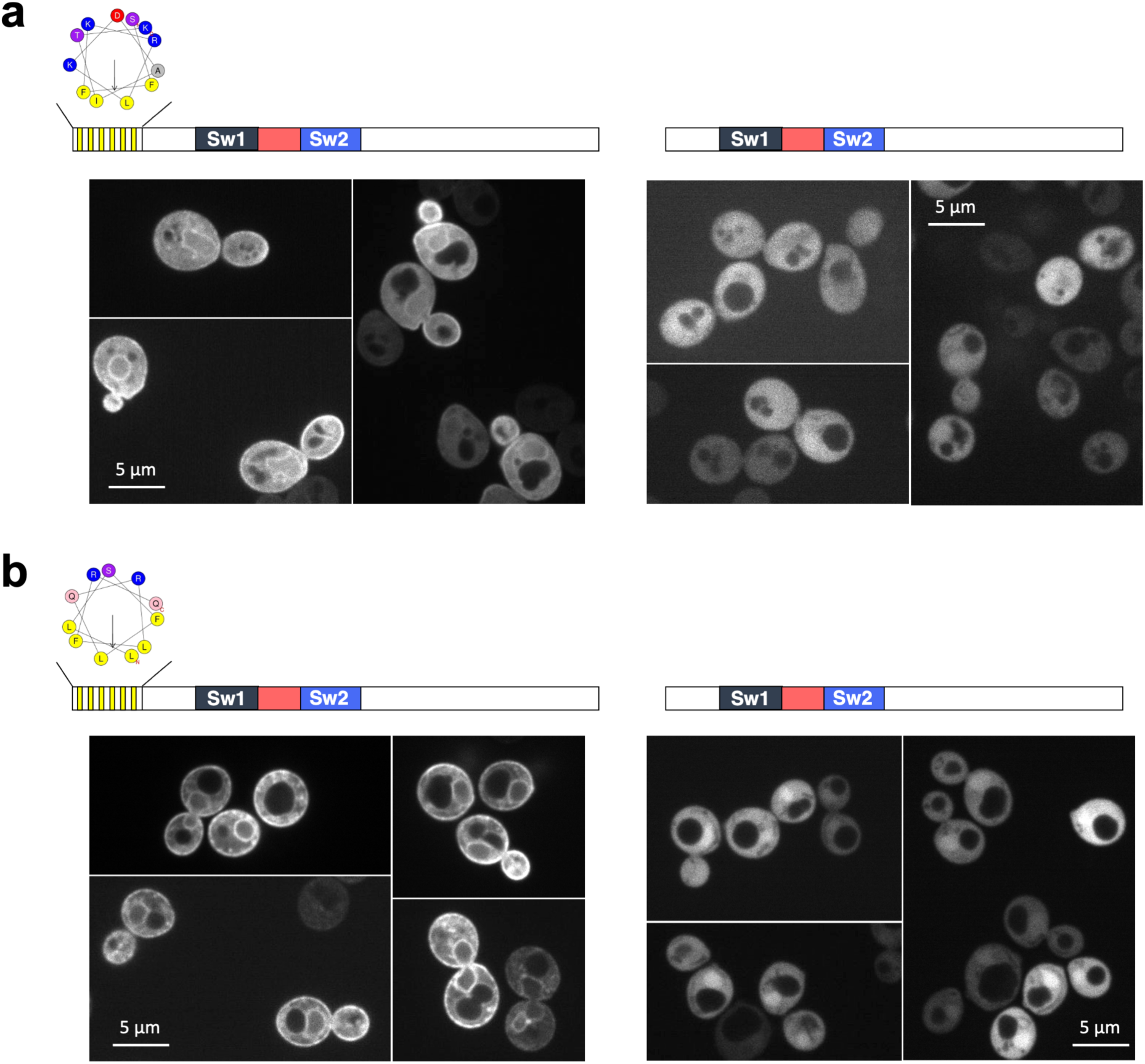
The N-terminal amphipathic helix (AH) of asgardarchaeote ArfR proteins is required for membrane localization in yeast cells. HodArfR1-Q72L (a) and wild type GerdArfR1 (b) were expressed as GFP fusion proteins in yeast cells. Representative images of full-length proteins (left panels) and N-terminal AH deletions (right panels) are shown. Schematic diagrams of the proteins expressed are shown above each set of microscopy images. Yellow and white vertical striped box represents the N-terminal AH, with Heliquest plot shown above (amino acids 2-13 for HodArfR1 (a); amino acids 2-11 for GerdArfR1 (b). Dark blue boxes, Switch I (Sw1); light blue boxes, Switch II (Sw2); red boxes, interswitch region.

The GerdArfR1-GFP protein localized to both the ER and PM in yeast cells, as was the case for HodArfR1-GFP, again indicating a lack of specificity for different eukaryotic membrane compartments (Fig. 3b, Fig. 4b, Extended Data Fig. 3). GerdArfR1-GFP had a higher membrane/cytosol ratio than HodArfR1-GFP when expressed in yeast cells, not only for the WT protein but also for the Q72L mutant (Fig. 3b, Extended Data Fig. 3). However, in contrast to HodArfR1, the GerdArfR1-Q69L mutant had the same, rather than an increased, level of membrane association compared to the GerdArfR1WT protein (Fig. 3b). As for HodArfR1, the predicted GTP-binding defective T27N mutation completely abolished ER and PM binding (Fig. 3b). These results support the conclusion that membrane binding of GerdArfR1in yeast cells requires that the protein be in its GTP-bound form. Deletion of the N-terminal AH of GerdArfR1 abolished localization to ER and plasma membranes in yeast cells, rendering the protein completely cytosolic (Fig. 3b, Fig. 4b), as was the case for HodArfR1. Together, these results support the conclusion that for these two asgardarchaeote ArfR proteins, membrane binding in cells is regulated by the guanine nucleotide bound, and is completely dependent on the N-terminal AH.

One of the two cultured Asgardarchaeota members (*Candidatus* Prometheoarchaeum syntrophicum, Lokiarchaeia) has been shown to have typical archaeal lipids, with gas chromatography–mass spectrometry detection of lipid species such as the isoprenoid C20-phytane^49^. The biophysical properties of membranes containing these methylated hydrophobic chains are similar to those containing diphytanoyl, which also has methylated hydrophobic tails^50^. Liposomes containing diphytanoyl are very permissive to adsorption of amphipathic helices^51,52^, suggesting that archaeal membranes would be ideal surfaces for an AH-mediated membrane binding mechanism. Indeed, AH binding to archaeal lipids has been demonstrated *in vitro* ^53^.

### Structural analysis of Asgard ArfR GTPases

To identify whether these selected asgardarchaeote ArfR proteins exhibit the unique structural properties of eukaryotic Arf family proteins, and in particular conformational changes upon GTP-hydrolysis that underpin membrane-localization by the AH, we determined the crystal structures of both their GTP– and GDP-bound forms. Because our cellular localization experiments indicate that the GTP-bound forms of both ArfR proteins are membrane bound in a manner dependent on their predicted N-terminal AHs, this portion was deleted, as is routinely done for eukaryotic Arf family proteins, to crystallise the GTP-bound form. Further, we mutated the catalytic glutamine residue to prevent possible GTP hydrolysis, as this is also routinely done for eukaryotic Arf proteins. Thus, the crystal structure of HodArfR1 bound to GTP (residues 15–184 with the Q72L mutation) was determined at 1.8 Å resolution (PDB entry 8OUK; statistics are given in Extended Data Table 1). HodArfR1-GTP possesses the typical fold of the Ras superfamily, with a central six-stranded-sheet (β1–β6) flanked by five-helices (α1–α5) (Fig. 5a, left). Well-defined electron density was observed for Switch I, interswitch and Switch II regions. The nucleotide-binding site reveals a classical organization with the magnesium ion possessing an octahedral coordination shell including two water molecules, two-phosphate Oβ and Oγ atoms, and side chains of Ser31 and Thr49 from the P-loop and Switch I, respectively (Fig. 5a, left). We next determined the crystal structure of the GTP-bound form of GerdArfR1 (PDB entry 8OUM; structural determination and analysis are given in Extended Data) which also possesses the typical fold of the Ras superfamily. The structural similarity between the GTP-bound forms of HodArfR1 and GerdArfR1 is high with a root-mean-square deviation (RMSD) value about 1.9 Å (based on 164 Cα-atoms). The structural similarity of both asgardarchaeote ArfR1 proteins is even better when compared with eukaryotic Arf proteins. For instance, superposition of human Arf1 with HodArfR1 and GerdArfR1 give a RMSD of 1.1 Å (161 Cα-atoms) for both comparisons, while that of Arf1 (PDB code 1O3Y) with Arf6 (PDB code 2J5X) gives a RMSD of 0.8 Å (162 Cα-atoms).

**Figure 5.**
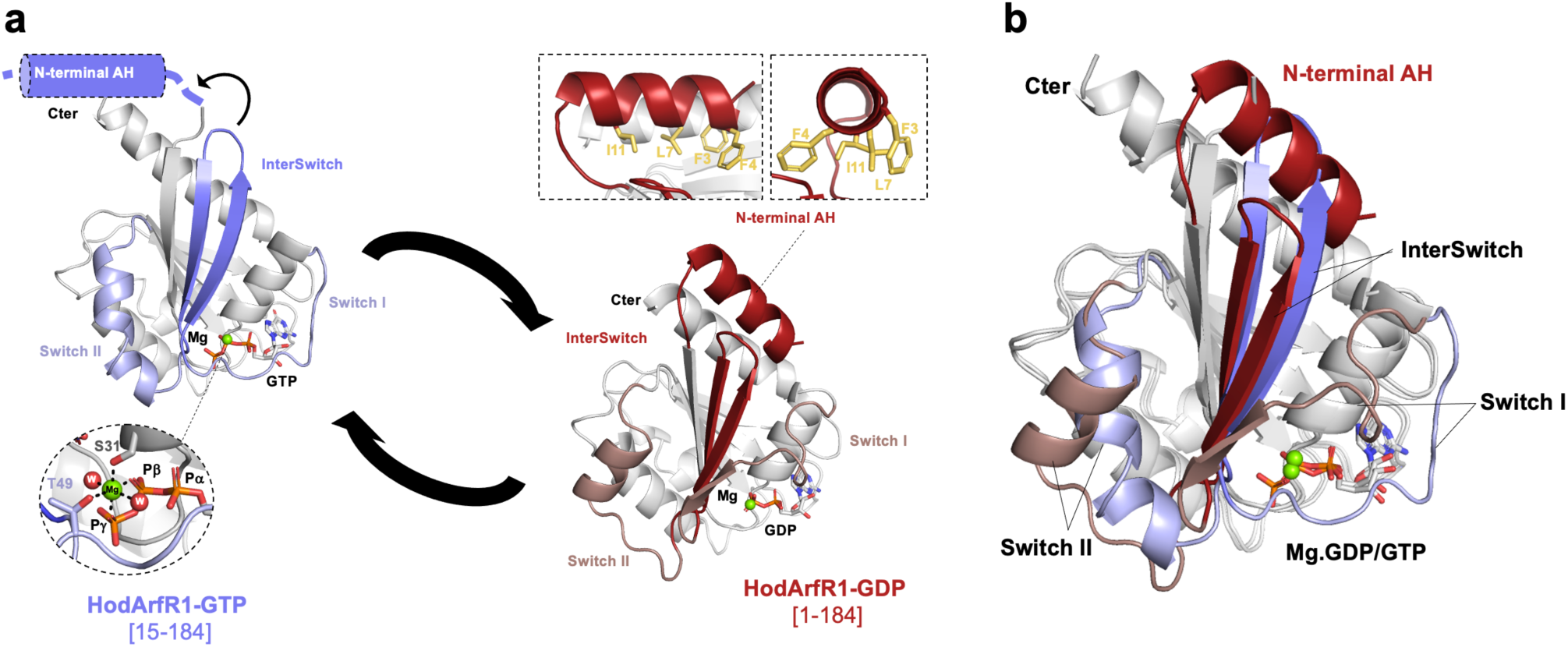
GDP/GTP structural cycle of HodArfR1. (a) Crystal structure of HodArf1-D14-Q72L bound to Mg.GTP (left panel). The protein is shown with the interswitch region in blue and the Switch I/Switch II regions in light blue. Below, the nucleotide binding site of the Mg.GTP is enlarged. The N-terminal helix, which is absent from the fragment crystallised, is shown schematically. Crystal structure of HodArfR1-FL-WT bound to Mg.GDP (right panel). The protein is shown with the N-terminal helix and the interswitch in red and the Switch I/Switch II in light red. Above, the N-terminal helix is enlarged and shown in two orientations, rotated by 90° with respect to each other; hydrophobic residues are indicated in yellow. Overall, the protein is shown with a cartoon representation, the magnesium ion is indicated by a green sphere and the GTP is depicted with a stick representation. (b) Superposition of the GTP-bound and GDP-bound forms of HodArf1. Colours are as described in part a.

We next determined the crystal structures of full-length wild type HodArfR1 and GerdArfR1 bound to GDP at 3.1 Å and 1.6 Å resolution, respectively (PDB entry 8OUL and 8OUN, respectively; statistics are given in Extended Data Table 1). For both, well-defined electron density was observed for the N-terminal AH, as well as for the switch regions. Overall, both HodArfR1-GDP and GerdArfR1-GDP possess the typical fold of the GDP-bound form of eukaryotic Arf proteins, with the N-terminal AH folded back on the core protein (Fig. 5a, right and Extended Data Fig. 5a, right). Hydrophobic residues from the N-terminal AH (for HodArfR1, Phe3, Phe4, Leu7, and Ile11) are buried in the hydrophobic pocket formed by residues from the tip of the interswitch region and the C-terminal helix (Fig. 5a, right; details about GerdArfR1-GDP structure are given in Extended data). Superposition of the GTP– and GDP-bound forms of HodArfR1 shows that switch regions, encompassing Switch I, interswitch and Switch II (Gln39 to Gly86) exhibit a drastically different conformation (Fig. 5b), in a way that parallels the structural rearrangements between GTP– and GDP-bound forms of eukaryotic Arfs. Indeed, the RMSD between the GTP– and the GDP-bound forms is 4.2 Å for the entire structure (169 Cα-atoms), while it decreases to 0.9 Å (121 Cα-atoms) when switch regions are removed from the calculation. Thus, the Switch I region (Gln39-Gly51) covers the nucleotide binding site in the GTP-bound form, while it adopts a conformation that fully uncovers the nucleotide binding site in the GDP-bound form. The C-terminal portion of Switch I adopts a β-strand secondary structure that makes main chain hydrogen bonds with the β2 of the interswitch region. In the GDP-bound form, the interswitch (Gly51-Gly71 consisting of the β2-β3 strands) is shifted toward the nucleotide binding site by a two-residue register adopting what we called hereafter the “retracted” conformation. In the GTP-bound form, the interswitch is “extended” and its β2-β3 hairpin (Asp60-Asn61) fully occupies the pocket where the N-terminal amphipathic helix lies in the GDP bound-form. Finally, Switch II (Gly71-Gly86), which forms two consecutive alpha helices in the GTP-bound form, adopts a unique helix, in the GDP-bound form. In the case of GerdArfR1, structural comparison of the GTP– and GDP-bound forms exhibit similar rearrangements in the switch regions (Extended Data Fig. 5b). Altogether, these results reveal that the unique features of the conformational switch between GTP– and GDP-bound forms of eukaryotic Arf family protein are conserved in these two asgardarchaeote ArfR proteins.

We directly compared the structures of the GDP-bound forms of HodArfR1 and GerdArfR1 with those of a selection eukaryotic Arf family proteins (Fig. 6). For this analysis, we identified three representative eukaryotic Arf family protein structures that were determined in the presence of their N-terminal AH: (i) the human Arf1 (named hereafter HsArf1; PDB code 1HUR^29^) which is a paradigm for eukaryotic Arf family proteins; (ii) the human Arl8A (HsArl8A; PDB code 4ILE; unpublished), which like HodArfR1 and GerdArfR1 lacks the myristoylation signal sequence and shares low sequence identity with HsArf1; and (iii) the unicellular parasite *Leishmania major* Arl1 (*Lm*Arl1; PDB code 2X77^54^) because of its evolutionary distance from mammalian Arf proteins. Overall, the structures are very similar in all pairwise comparisons with the interswitch region in a retracted position and the N-terminal AH folded back between the interswitch tip and the C-terminal helix (Fig. 6 and Extended Data Fig.5). Further, Switch I adopts a β-strand secondary structure that interacts with the β2-strand of the interswitch uncovering the GDP nucleotide binding site and Switch II, which are more or less flexible and exhibit helical segments at this C-terminus. But locally, some variations are observed between these structures, especially in the exact position/orientation of the N-terminal AH (Fig. 6a-c and Extended Data Fig.5). Such a difference is not unexpected considering that the interactions between the AH and the rest of the protein are mainly hydrophobic, which allow non-specific interactions and thus sliding movement. Also, the difference in length and in the primary sequence of the N-terminal AH (Fig. 5d), especially the number, position and nature of the hydrophobic residues account for such variations. Thus, pairwise comparisons of full Arf protein structures give RMSDs (using Cα atoms) ranging from 1.5 Å to 4.0 Å and decreasing from 0.9 Å to 1.9 Å when N-terminal AH and Switch I/Switch II sequences are removed (noted below as RMSD 1.5-4.0/0.9-1.9 Å; Extended Data Fig.5). HsArf1 and *Lm*Arl1 are structurally very close (RMSD 1.5/0.9 Å), while the difference between human Arf1 and Arl8A is greater (RMSD 2.9/1.3 Å), but similar to that observed between archaeal HodArfR1 and the three eukaryotic Arf family protein structures (2.6-2.8/1.2-1.3 Å) or between archaeal GerdArfR1 and HsArf1/LmArl1 structures (RMSD 2.8-3.0/1.4-1.7 Å). One difference comes from pairwise comparison between archaeal GerdArfR1 and HsArl8 that exhibit greater RMSDs of 3.6/1.9 Å, which are similar to what was measured between HodArfR1 and GerdArfR1 (RMSD 4.0/1.9 Å). Thus, these structural analyses reveal that differences found between archaeal and eukaryotic Arf family proteins were no greater than those found between different pairs of eukaryotic Arf family proteins (for instance HsArf1 vs HsArl8A) or between archaeal HodArfR1 and GerdArfR1 proteins themselves (Extended Data Fig. 5).

**Figure 6.**
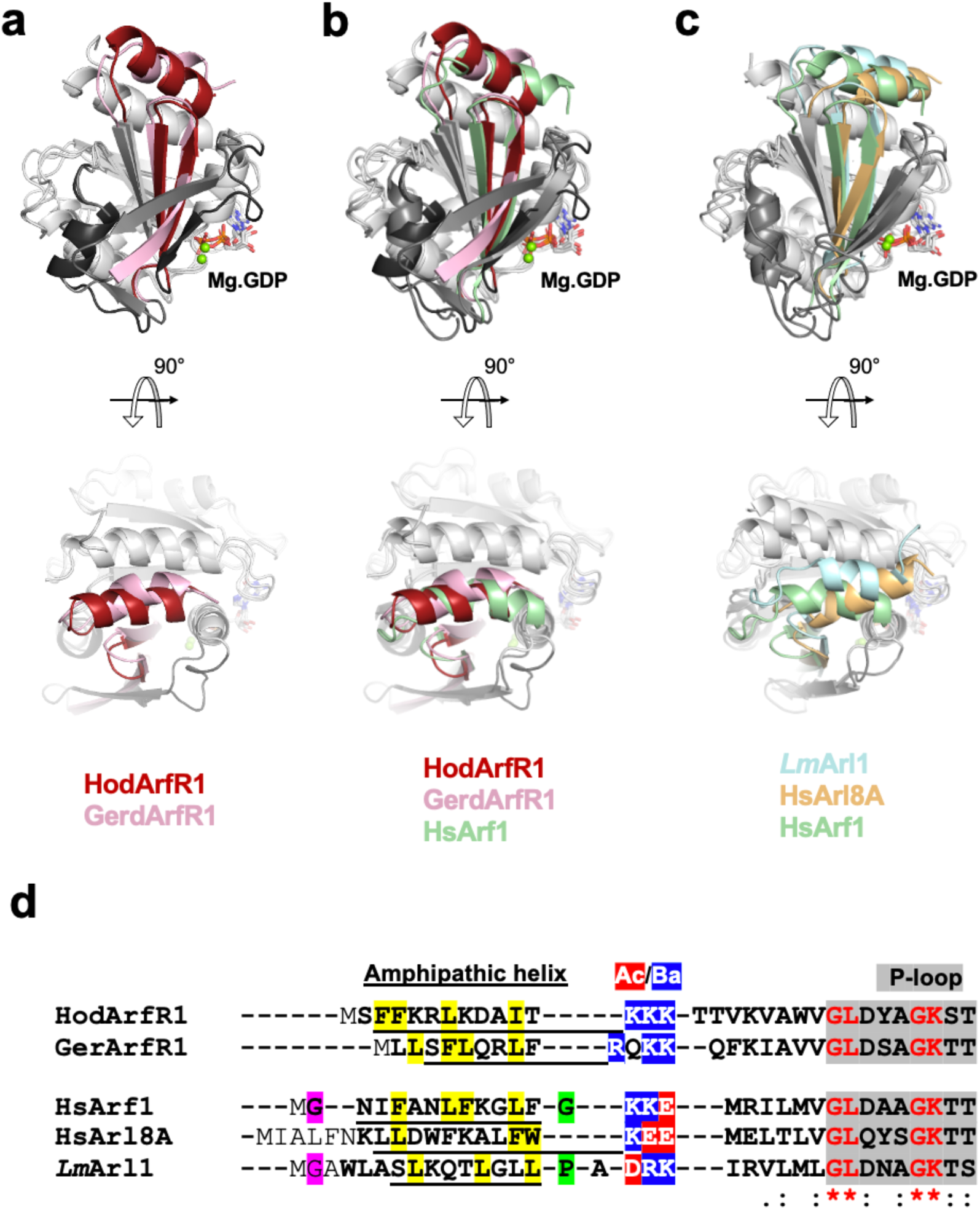
Structural comparison of the GDP-bound form of asgardarchaeote and eukaryotic Arf family proteins. (a) HodArfR1-GDP is compared to GerdArfR1-GDP. HsArf1-GDP is compared to HodArf1-GDP and GerdArfR1-GDP (b) and to HsArl8-GDP and *Lm*Arl1-GDP (c). Above, face orientation showing the N-terminal and the interswitch regions in colours and Switch I/Switch II regions in grey; below, a 90° top rotation focusing on the N-terminal AH. Superposition was performed on residues of the P-loop, strands β1-β3 and the C-terminal helix. The same orientation is shown for all superpositions. Structural superposition of these Arf proteins was performed on residues from the P-loop, the interswitch, the β4 and the C-terminal helix which altogether allow the comparison of the position of the N-terminal AH and the switch regions relative to the rest of the core protein. (d) Structure-based sequence alignment of HodArfR1 and GerdArfR1 with three eukaryotic Arf proteins (human Arf1, human Arl8A and *Leishmania major* Arl1). Bold letters indicate residues present in the structure; normal text, amino acids that are not present in the structure (either due to absence in the plasmid or without electronic density (flexible). Hydrophobic amino acids from the AH and the glycine at position 2 are highlighted in yellow and pink, respectively.

Altogether, our experimental structural studies clearly show that HodArfR1 and GerdArfR1, and probably other asgardarchaeote ArfR proteins, share with eukaryotic Arf family proteins the interswitch toggle that dislodges the N-terminal AH, which is the basis of the regulation of cytosol-membrane localization of these proteins.

## Conclusions

Here we report a newly delineated group of prokaryotic Ras superfamily GTPases, ArfR. These proteins are found prominently amongst the Asgardarchaeota, the only exception being a probable horizontal transfer of ArfR into a single Thermoproteota (TACK) lineage. The ArfR GTPases most likely evolved in the common ancestor of the Asgardarchaeota, and strikingly, are found in the lineages most closely related to eukaryotes. Crucially, the ArfR group is clearly from where the eukaryotic Arf family GTPases arise. We have also demonstrated using experimental cellular and structural approaches that membrane-localization *via* the N-terminal AH, a hallmark feature of the Arf family, is conserved in the ArfR representatives that we tested, strongly supporting direct inclusion of ArfR in the Arf family. These results pinpoint the origin and timing of acquisition of an important protein family in the last Asgardarchaeota and eukaryote common ancestor, with profound implications for eukaryogenesis. At a minimum, any scenarios for eukaryogenesis must include an Arf GTPase in the archaeal contributor that has the capacity to localize to membranes, via guanine nucleotide regulation of AH membrane binding.

A regulated actin cytoskeleton has been demonstrated experimentally in Asgardarchaeota using multiple approaches^16–19,55^. Tubulin has also been shown experimentally to be present in Asgardarchaeota^20^. ESCRT complexes, which are involved in endocytic membrane trafficking in eukaryotes, have also been demonstrated to exist in the Asgardarchaeota^14,56,57^. However, the ESCRT machinery mediates budding of vesicles into the lumen of organelles and towards the exterior of the cell, not towards the cytoplasm as would be required for the establishment of a membrane trafficking system. Although the number of identified ESGs related to membrane trafficking function encoded in asgardarchaeote genomes continues to increase^9^, experimental evidence that they carry out these functions is scarce. Among proteins required for cytoplasmic vesicle formation and fusion, the closest demonstration of functional homology is the capacity of the Hodarchaeales and Heimdallarchaeaceae SNARE homologs to form complexes *in vitro*^15^. So whereas the presence of membrane-trafficking ESGs is tantalizing, the implications are still somewhat nebulous. Our data identifies a protein family in Asgardarchaeota whose eukaryotic members include those with key functions in eukaryotic membrane trafficking, with experimental support for the conservation of their hallmark mechanism of membrane localization. Based on these results, it seems that the archaeal contributor to eukaryogenesis was poised for cytoskeleton-modulated endomembrane function, prior to the mitochondrial acquisition.

## Acknowledgements

We especially thank Drs. B. Gigant and V. Campanacci for help with X-ray data collection and scaling, and C. Besse and M. Chenon for plasmid construction. We are most grateful to the machine (SOLEIL synchrotron, Saint-Aubin, France) and beamline groups (PROXIMA 1 and PROXIMA 2A) for making these experiments possible. This work has benefited from the crystallization (S. Plancqueel and A. Vigouroux) and protein-protein interaction (M. Aumont-Niçaise) platforms of I2BC supported by French Infrastructure for Integrated Structural Biology (FRISBI) ANR-10-INBS-05. We thank Dr. V. Campanacci for careful reading and comments on the manuscript. This study was supported by the CNRS, France and grant ANR-20-CE13-0007 from the ANR, France, to JM and CLJ, and the LERCO project number CZ.10.03.01/00/22_003/0000003 via the Operational Programme Just Transition to ME.

## Author contributions

RV, RC, MA, CC and KT contributed to the acquisition and analysis of data; PL contributed to data analysis; ME, JM, JBD and CLJ contributed to the conception and design of the work, data analysis and writing the manuscript.

## Materials & Correspondence

Correspondence and requests for materials should be addressed to Marek Elias; Julie Ménétrey; Joel B. Dacks; or Catherine L. Jackson.

## Methods

### Sequence searches, annotation, and analyses

Genes of interest in target genomes (most of them in the form of MAGs) were identified with BLAST^58^, using a stand-alone version or the on-line implementation at the NCBI server https://blast.ncbi.nlm.nih.gov/Blast.cgi; default parameters). For identification of candidate Ras superfamily sequences, various eukaryotic and prokaryotic protein sequences representing different major subgroups of the Ras superfamily were used as queries in the searches. In the case of MAGs without a gene prediction in the respective database, GetORF (https://www.bioinformatics.nl/cgi-bin/emboss/getorf) with default settings was used for generating a working set of protein sequence prediction. Predicted protein sequences sets were searched with BLASTp, but all genome assemblies were additionally checked with tBLASTn to identify genes that had escaped annotation. Hits with e-value lower than 1.0 were considered as candidate sequences. Whether a candidate sequence is a member of the Ras superfamily was determined based on the results of an online BLASTp searches on the NCBI website against the “non-redundant protein sequences” (nr) database, and of a CD-Search^59^ against the Conserved Domain Database (https://www.ncbi.nlm.nih.gov/Structure/cdd/wrpsb.cgi^60^). Protein sequences that according to the CD-Search contained a GTPase domain matching as best hits GTPase domains not considered to correspond to the Ras superfamily (as delimited by Leipe et al.^61^), were discarded. Further sequences were discarded if sharing the highest sequence similarity (as scored by BLASTp) with proteins other than members of the Ras superfamily, except for the cases of multidomain proteins containing another protein domain along with a Ras superfamily GTPase domain. We also retained proteins without a significant CD-Search match to any recognisable protein domain, but with a high sequence similarity (as determined by BLASTp) to other candidate Ras superfamily proteins from Asgardarchaeota identified as described above.

In the earliest phase of the project we tried to identify and annotate Ras superfamily GTPases in every asgardarchaeote genome reported in the literature^7,8,41,62^, but a comprehensive analysis of Ras superfamily genes in Asgardarchaeota became unfeasible with the dramatic increase in the rate new asgardarchaeote MAGs have started to accumulate in the literature^9,10,63–67^ and databases since 2019. To account for this development, as well as to cope with the incongruent taxonomy asgardarchaeote lineages proposed in different studies, the highly varying quality of asgardarchaeote MAGs (concerning their completeness and the level of contamination), and frequent mistakes in the taxonomic assignment of MAGs deposited in the NCBI database, we performed our own survey of archaeal genome assemblies (available as of May 2022) to delineate the main phylogenetic lineages of Asgardarchaeota represented by the assemblies. To this end we primarily used the sequence of RNA polymerase subunit B (RPOB) encoded by archaeal genomes as a phylogenetic marker, which proved to provide results highly congruent with multigene analyses of the asgardarchaeote phylogeny (Fig. S1). Based on these analyses, and consistent with the most recent published reassessment of the asgardarchaeote phylogeny^9^, “Freyrarchaeota”^67^ and “Freyarchaeota” (NCBI taxonomy database, txid2827219) are considered synonymous with Jordarchaeia^66^, and “Borrarchaeota”^10^ are considered synonymous to Sifarchaeia^66^. In addition, we follow Eme et al.^9^ by accepting the revised ranking of the originally proposed separate phyla of the “Asgard superphylum” as classes, orders or families, and subsume the putative phylum “Sigynarchaota”^67^ into Lokiarchaeales based on the results of our phylogenetic analysis. They also pointed to the existence of a novel asgardarchaeote lineage that in the meantime has been recognized as the class Asgardarchaeia^9^. Finally, for the asgardarchaeote lineage originally known as “ACLG”^66^ we adopt the name Atabeyarchaeia proposed in a recent preprint^68^. Guided by the RPOB phylogeny, additionally aided by a tree inferred from an alternative phylogenetic marker DNA polymerase I to allow for placement of a few MAGs missing the RPOB gene (not shown), we selected for further analyses of the Ras superfamily genes a phylogenetically representative subset of 62 genome assemblies (Table S2), considering the following criteria: 1) to include at least one representative for every major asgardarchaeote lineage named in the literature; 2) to sample more or less evenly across the phylogenetic diversity of each major group; 3) to include prominent representatives, such as the initially reported MAGs previously analysed in detail, the completely assembled (closed) MAGs, and the genome of the first cultivated asgradarchaeote (*Candidatus* Prometheoarchaeum syntrophicum^49^, to prioritize higher-quality MAGs (based on the quality estimates reported for the MAGs in the literature).

Following the procedure described above, our analysis of the 62 asgardarchaeote genome assemblies yielded a dataset composed of >2900 Ras superfamily proteins (Table S3). Six proteins turned out to be composed of two different Ras superfamily GTPase domains; these were split and each domain treated separately in subsequent analyses (Table S3, column “Note(s)”). For further analyses (except for the analysis of the presence of an N-terminal AH, see below), all sequences were trimmed to GTPase domain to avoid biases introduced by the extra domains present in some of the proteins. Phylogenetic analyses were used to delineate major subgroups of asgardarchaote Ras superfamily proteins, to illuminate their phylogenetic relationship to eukaryotic GTPases, and to annotate individual sequences. Generally, sequence alignments were built with MAFFT version 7^69^ with default settings, manually checked and corrected whenever necessary. In the case of the for phylogenetic analysis displayed in Extended Fig. 1, the alignment was constructed using PASTA^70^. Alignments were trimmed by a stand-alone version of trimAl^71^ using the –gappyout option. Maximum likelihood (ML) phylogenetic trees were calculating using IQ-TREE^72^ with the substitution model selected by ModelFinder implemented in IQ-TREE. Branch supports were assessed by the SH-aLRT test^73^ and the ultrafast bootstrap approximation^74^ are also part of IQ-TREE. For some analyses, the phylogenetic pipeline provided by the ETE3 server (https://www.genome.jp/tools-bin/ete) was used. A number of preliminary analyses were performed to get initial insights the diversity of asgardarchaote Ras superfamily proteins and evaluate the robustness of the results depending on sequence selection. The details of final analyses included here to document the key findings are summarized in Table S4 and mentioned also in the legends to the respective figures.

All Ras superfamily sequences identified in the first asgardarchaeote MAGs reported^7,8,41,62^, were combined with the GTPase dataset analysed by Klinger et al.^22^ comprising both eukaryotic and prokaryotic sequences (including also sequences from Lokiarchaeota archaeon GC14_75). The tree inferred from this dataset (Extended Data Fig. 1) enabled us to define ArfR and Rup3 as novel asgardarchaeote Ras superfamily subgroups added to those distinguished before. Ras superfamily sequences from the remaining asgardarchaote genome assemblies were then annotated (i.e., assigned to a particular predefined subgroup) by combining them with a representative subset (“annotation dataset”) of 191 sequences from the aforementioned dataset, and inferring a tree from the new alignment (created *de novo* with MAFFT). This annotation procedure was applied either separately to all Ras superfamily proteins encoded by a single asgardarchaeote MAG, or sequences from multiple related MAGs were included in the same analysis together if their number encoded by the given genomes was low. Newly analysed sequences were assigned to a particular Ras superfamily subgroup if they were part of a clade with the respective reference sequences, regardless of the statistical support for the monophyly of the clade. This approach left 159 sequences unannotated, but 35 of them were further tentatively annotated based on their comparison to the whole set of asgardarchaeote Ras superfamily with BLASTp. The remaining unannotated sequences represent mixture of rapidly evolving derivatives (possibly even pseudogenes) of the recognized GTPase subgroups and potentially novel GTPase subgroups conserved in particular taxa that are yet to be investigated in detail before they are formally defined. All phylogenetic trees were manually checked and sequences that resided on obviously long branches were labelled as divergent in Table S3, while sequences with GTPase domain shorter than 100 amino acid residues were flagged as incomplete in Table S3. The so-called “representative dataset” composed of 2626 sequences was created by combining two reference korarchaeial sequences annotated as ArfR (WP_083758176.1 and WAOH01000054.1) with the set of asgardarchaeote Ras superfamily sequences excluding the incomplete ones, those flagged as divergent, those that remained unannotated or were annotated only based on BLASTp searches, and those representing the Rup3 group (having the GTPase modified by unusual insertions; Fig. S2B). A tree inferred from the “representative dataset” (Fig. S3) confirmed the annotation of the sequences included.

To allow for more thorough phylogenetic analyses, the “representative dataset” was subjected to a ScrollSaw procedure, originally designed for a phylogenetic analysis of the Rab GTPase family^43^ and since then used in similar contexts^23,75^. Here the principle is to reduce the complexity of a sequence dataset by concentrating on more slowly evolving sequences with readily identifiable orthologs across organismal lineages. To this end, the asgardarchaeote sequences were divided into 17 groups corresponding to major asgardarchaeote taxonomic groups (putative phyla) recognized at that time (considering “Sigynarchaeota” as a separate group; Table S2). Alignments were created for sequences from each possible pair of 17 taxonomic groups (136 alignments) and genetic distances were estimated using the ML method by the Tree-Puzzle 5.3^76^ with the WAG+Γ+I substitution model. A custom Python script was then used to identify all so-called minimal-distance pairs for each resulting distance matrix, yielding a set of 1341 sequences, which was further reduced by removing those sequences that were part of only one minimal-distance pair, resulting in a set of 777 asgardarchaeote Ras superfamily sequences. We then added to them sequences of the two proteins (OLS23013.1 and NHJ47629.1) that were targeted by our experimental work but which did not pass the filtering by the ScrollSaw approach, and the two aforementioned representative korarchaeial ArfR sequences. On the other hand, two related asgardarchaeote sequences (WAMJ01000037.1_559 and OLS29018.1), both clearly annotated as ArfR (see Fig. S3), were eventually removed from the ScrollSaw-derived dataset, as they for unknown reasons behaved as “rogue” sequences, exhibiting an unstable position in preliminary analyses and decreasing statistical support of major Ras superfamily groups. The resulting so-called “final ScrollSaw dataset” composed of 779 sequences was then combined for a phylogenetic analysis (Fig. 1; Fig. S4) with reference sequences representing the prokaryotic Rup1, Rup2, MglA1, and MglA2-5 subgroups and the different eukaryotic subgroups of the Ras superfamily, selected on the basis of previous analyses^23,43^. Given the special focus on the relationship of ArfR and eukaryotic Arf family GTPases, sequences representative for all Arf family paralogs found in the LECA (including SRβ) were included.

For a dedicated analysis of the Rup3 group (Fig. S2), a comprehensive search of archaeal genome assemblies available in the NCBI database was performed with BLAST, with the additional Rup3 sequences (those not encoded by any of the 62 genome assemblies selected for comprehensive analyses; Table S5) recognized based on the characteristic insertions in the GTPase domain and added to the analysis. Similarly, a comprehensive search for ArfR genes in korarchaeial genome assemblies available in the NCBI database was performed with BLAST for a dedicated phylogenetic analysis. Searched were all assemblies identified as representing Korarcheia based on a phylogenetic analysis of the RPOB protein (Fig. S5), which enabled us to recognize as korarchaeial a number of assemblies not classified as such in the NCBI database. All identified korarchaeial ArfR sequences are listed in Table S5 (and were included in the phylogenetic analysis presented as Fig. S6). The occurrence of Ras superfamily genes potentially related to the eukaryotic Arf family in archaeal genome assemblies in the NCBI database not assigned to Asgardarchaeota or Korarchaeia was investigated by tBLASTn searches with the human Arf1 sequence as a query. The individual hits were analysed by backward BLAST searches against the NCBI nr database and our custom GTPase sequence database, and the taxonomic assignment or the respective assemblies was scrutinized by determining the phylogenetic position of the RPOB or DNA polymerase I proteins encoded by the assemblies. All candidates were found to represent misidentified asgardarchaeote, korarchaeial, or eukaryotic sequences (Table S1).

The sequence similarity-based clustering analysis (Extended Data Fig. 2) was performed using CLANS^77^. 2994 sequences combining the “representative dataset” (see above), Rup1, Rup2 and Rup3 sequences, and a selection of eukaryotic Ras superfamily GTPases (those used in the phylogenetic analysis shown as Fig. S4) were subjected to all pairwise comparisons with BLASTp using the online utility included in the MPI Bioinformatics toolkit (https://toolkit.tuebingen.mpg.de/tools/clans); keeping the default parameter setting. The resulting matrix was used as an input for the CLANS executable, which was run with the default settings until no major changes in the visible clustering pattern were observed. As the clustering procedure is non-deterministic, several independent runs were performed and confirmed that the general clustering pattern is robust. CHROMA ver. 1.0^78^ was used for annotation of the Rup3 sequence alignment shown in Fig. S2B. To evaluate the presence of an N-terminal AH in ArfR sequences, those with validated complete N-terminal regions were further sorted according to the presence of additional domains upstream or downstream of the G domain. One hundred ArfR sequences (160-220 amino acids long) with no large domains flanking the G domain were screened using Heliquest^79^ for the presence of a predicted membrane-binding N-terminal AH. AHs with the capacity to bind membranes can have a range of chemical properties^80^, so we chose parameters that allowed for a diversity of such AHs to be identified. Forty-five ArfR proteins were identified with a predicted membrane-binding AH at the N-terminus, including HodArfR1 and GerdArfR1.

### Yeast strains and microscopy

*Saccharomyces cerevisiae* strains BY4742 *MATα his3Δ1 leu2Δ0 lys2Δ0 ura3Δ0* and EY0987 Sec13::mRFP *MATα his3Δ1 leu2Δ0 lys2Δ0 ura3Δ0 Sec13::mRFP-kanMX6* were used in this study. Yeast plasmids used in this study are listed in Extended Data Table 3. pRS416-prADH-GFP (pCLJ1101) was a gift from Sebastien Leon, Institut Jacques Monod, Paris, France and pHCMM4 was a gift from Hugo Cesar Medina-Munoz and Marvin Wickens (Addgene plasmid #170359; RRID:Addgene_170359)^81^. Yeast media and lithium acetate plasmid transformations were as described previously^80^.

Synthetic genes encoding HodArfR1 (GenBank: OLS23013.1) and GerdArfR1 (GenBank: NHJ47629.1), and optimized for expression in yeast (Eurofins Genomics Europe, Ebersberg Germany), were cloned into pCLJ1101. Mutants were obtained either by site-directed mutagenesis or by synthetic gene synthesis. Yeast strains transformed with expression plasmids were grown to mid-logarithmic phase in selective medium, then imaged. Images were acquired at room temperature with spinning-disk confocal microscope CSU-W1 (Yokogawa – Andor) driven by MetaMorph or ZEN 2 software. 28 z-sections separated by 0.26 µm were acquired for each field of cells using excitation/emission wavelengths 488/509 and 545/572. Localization of GFP-tagged proteins was analysed in ImageJ by selecting an appropriate z-section, choosing cells with well-defined organelle localization, and drawing a line across the cell (examples are shown in Fig. 3a, b, yellow lines). Fluorescence intensity was obtained for each pixel along the line, and peaks at either the cell boundary (plasma membrane) or ER were determined. Plots of pixel intensities across the yellow lines in the left panels of Fig. 3 are shown in the middle panels of Fig. 3a, b. The average cytoplasmic fluorescence intensity was obtained by selection of a portion of the line passing over cytoplasm free of organelles, and averaging the fluorescence values obtained. The membrane/cytosol ratio was calculated by dividing the membrane fluorescence value by the average cytosolic fluorescence value. For each transformed strain, 15-30 cells, from separate fields, were analysed, and the mean and standard deviation of the calculated membrane/cytosol ratios was obtained using Graphpad Prism software (version 10.2.0). At least 3 independent experiments were performed for each transformant.

### Structural studies of the archaeal HodArfR1 protein

The *Escherichia coli* optimised cDNAs coding for HodArfR1-FL-WT (aa. 1-184) and HodArfR1-D14-Q72L (aa. 15-184) (GenBank: OLS23013.1) were obtained by gene synthesis. Both HodArfR1-FL-WT and HodArfR1-D14-Q72L inserts were subcloned into the pET21a plasmid and produced in *E. coli* BL21 DE3 as an uncleavable C-terminal His6 tag fusion proteins. Cells were induced with 0.3 mM IPTG, 4 h at 37°C and then centrifuged at 4 000 rpm for 30 min at 4°C. Pellets were suspended in a solution containing 50 mM Tris pH 8.0, 500 mM NaCl, 1 mM DTT, 1 mM MgCl_2_, 20 mM Imidazole pH 8.0, benzonase and anti-proteases. After cell disruption by sonication, the lysate was ultracentrifuged at 22 000 g for 35 min at 4°C and the supernatant was filtered on 0.22 mm filters and then loaded on a His-Trap 5mL column (GE Healthcare) equilibrated with 50 mM Tris pH 8.0, 500 mM NaCl, 1 mM MgCl_2_ 1 mM DTT, 20 mM imidazole pH 8.0 and 10 μM GDP (for HodArfR1-FL-WT) or GTP (HodArfR1-ι114-Q72L). Protein elution was performed in the same buffer containing 500 mM imidazole pH 8.0. Finally, all proteins were further purified by gel filtration on a HiLoad 16/60 Superdex 75 column (GE Healthcare) in 40 mM Tris pH 8.0, 200 mM NaCl, 1 mM MgCl_2_, 1 mM DTT and 10 μM GDP or GTP. Proteins were stored at –80°C after addition of 1 mM GTP or 1 mM GDP in the storage buffer for HodArfR1-ι114-Q72L and HodArfR1-FL-WT, respectively.

Crystals of HodArfR1-ι114-Q72L bound to Mg.GTP were obtained by the vapour diffusion method at 290 K in sitting drops using equal amounts of protein at 15.6 mg ml^-1^ and a reservoir solution consisting of 2.0 M sodium formate and 0.1 M HEPES pH 7.5. Crystals were transferred for a few seconds to a cryo-protectant composed of a reservoir solution supplemented with 25% glycerol and frozen in liquid nitrogen. Diffraction data were collected at 100 K on the PROXIMA-2 beamline (SOLEIL Synchrotron). Crystals diffracted up to 1.8 Å and belonged to the tetragonal space group P4_3_2_1_2 with one molecule in the asymmetric unit. X-ray data were integrated and scaled using XDS^82^. The structure was determined by molecular replacement with PHASER^83^ using as a search model the homology model of HodArfR1-D14-Q72L-GTP generated using Phyre2^84^. AutoBuild^85^ was initially used to automatically refine the structure, which was followed by final iterative refinements using buster with the TLS option^86^ and graphical building performed using COOT^87^. The electron density clearly allowed the tracing of the complete protein from the Met14 to the Ile184 of HodArfR1-D14-Q72L. The GTP molecule is clearly defined in the electron density, as well as the Mg^2+^ ion and its two coordinated H_2_O molecules.

For the crystallisation assays with the GDP-bound form of HodArfR1-FL-WT, nucleotide exchange was performed using purified protein incubated in 2.5 mM EDTA for 5 min at room temperature in the presence of 10× molar excess GDP (5 mM GDP). The exchange was terminated by addition of 5 mM MgCl_2_ and excess nucleotide was removed by dialysis. Crystals of HodArfR1-FL-WT bound to Mg.GDP were obtained by the vapour diffusion method at 290 K in sitting drops using equal amounts of protein at 6.5 mg ml^-1^ and a reservoir solution consisting of 30% PEG 4000, 0.2 M lithium sulfate, 0.1 M Tris-HCl pH 8.5 (condition 89 of the QUIAGEN Classic Suite screen). For data collection, crystals were transferred for a few seconds to a cryo-protectant composed of a reservoir solution supplemented with 18% ethylene glycol and frozen in liquid nitrogen. Diffraction data were collected at 100 K on the PROXIMA-1 beamline (SOLEIL Synchrotron). Crystals diffracted up to 3.1 Å and belonged to the tetragonal space group P4_3_2_1_2 with three molecules in the asymmetric unit. X-ray data were integrated and scaled using XDS^82^ and anisotropy correction of the raw data set was performed using STARANISO^88^. The structure was determined by molecular replacement with PHASER^83^ using as a search model the HodArfR1-Δ14-Q72L-GTP structure (this study) lacking the Switch I-interswitch-Switch II region (residues 40-88). Manual building was used to dock the N-terminal helix and the interswitch in the initial extra electron density, and then to build the Switch I and Switch II region in the first refined extra electron density. Refinement was then carried out using buster with the autoNCS option^86^ and the graphical building was performed using COOT^87^. The GDP molecule and the Mg^2+^ ion are clearly defined in the electron density. The three molecules (A, B and C) in the asymmetrical unit are virtually identical, except in the Switch II region that exhibits slight differences due to crystal packing variations. Thus, their entire Cα traces show RMSDs ranging from 0.28 Å to 0.40 Å (calculated over 177-180 residues) and ranging from 0.20 Å and 0.24 Å (when the Switch II residues 69-78 is removed).

Data collection and refinement statistics for HodArfR1-Δ14-Q72L-GTP (PDB code 8OUK) and HodArfR1-FL-WT-GDP (PDB code 8OUL) structures are presented in Extended Data Table 1. Structural analysis and pairwise superposition of Arf proteins were performed using Pymol software^89^. Figures were produced using the same software. Cloning, protein expression and purification for GerdArfR1 proteins, as well as crystallisation conditions, data collection, and structure determination are presented in Extended Data Methods.

## Extended Data

### Extended Data Methods

#### Structural study of the archaeal GerdArfR1 protein

Cloning of GerdArfR1-FL-WT (residues 1-181) and GerdArfR1-Δ12-Q69L (residues 13-181) (GenBank: NHJ47629.1), as well as protein expression in *E. coli*, purification and storage were identical to those carried out for HodArfR1 protein (Methods, main text).

Crystals of the GerdArfR1-Δ12-Q69L bound to Mg.GTP were obtained by the vapor diffusion method at 290 K in sitting drops using equal amounts of protein at 11.8 mg ml^-1^ and reservoir solution consisting of 8.5% PEG 8000, 0.1M MgCl2 and 0.1M Tris pH 7.0. Crystals were transferred briefly to a cryo-protectant composed of reservoir solution supplemented with 25 % glycerol and frozen in liquid nitrogen. Diffraction data were collected at 100 K on the PROXIMA-2 beamline at the SOLEIL Synchrotron. Crystals diffracted up to 2.7 Å and belonged to the monoclinic space group P2_1_ with five molecules in the asymmetric unit. X-ray data were integrated and scaled using XDS^1^. The structure was determined by molecular replacement with PHASER^2^ using as a search model a homology model of the GTP-bound form of GerdArfR1-D12-Q69L generated using AlphaFold2^3^. Refinement was carried out using Buster with TLS and NCS options^4^ and the graphical building was performed using COOT^5^. The polypeptide chain of the five molecules shares clear electron density from Lys13 to Lys180. In all molecules, the GTP molecule, as well as the magnesium ion and its coordinated water molecules are clearly defined in the electron density. All five molecules are virtually identical with an RMSD ranging from 0.23 to 0.37 Å on their entire Cα traces (calculated over residues 167-173).

For the crystallization assays with the GDP-bound form of GerdArfR1-FL-WT, nucleotide exchange was performed using purified protein incubated in 2.5 mM EDTA for 5 minutes at room temperature in the presence of 10X molar excess GDP (5 mM GDP). The exchange was terminated by addition of 5 mM MgCl_2_ and excess nucleotide was removed by dialysis. Crystal of the GerdArfR1-FL-WT bound to Mg.GDP were obtained by the vapour diffusion method at 290 K in sitting drops using equal amounts of protein at 8.3 mg ml^-1^ and reservoir solution consisting of 18% PEG 8000, 0.2 M calcium acetate, 0.1M sodium cacodylate pH 6.5 (condition 67 of the QUIAGEN Classic Suite screen). For data collection, crystals were transferred briefly to a cryo-protectant composed of reservoir solution supplemented with 25 % glycerol and frozen in liquid nitrogen. Diffraction data were collected at 100 K on the PROXIMA-2 beamline at the SOLEIL Synchrotron. Crystals diffracted up to 1.65 Å and belonged to the monoclinic space group P2_1_ with two molecules in the asymmetric unit. X-ray data were integrated and scaled using autoPROC^6^. The structure was determined by molecular replacement with PHASER^2^ using as a search model a homology model of GerdArfR1-FL-WT-GDP generated using Phyre2^7^, lacking the amphipathic N-terminal helix, Switch I and Switch II regions, but keeping the retracted interswitch region. Automatic building was performed using Arp/Warp^8^ and then iterative refinement was carried out using Buster with TLS option^4^ and the graphical building using COOT^5^. The polypeptide chain was modelled in clear electron density from Leu2 to Leu181. In both molecules A and B, the GDP molecule, as well as the magnesium ion and its coordinated water molecules were clearly defined in the electron density. Molecules A and B are virtually identical, but slight movements are observed for the Switch I-interswitch-Switch II region between the two molecules due to different crystal packing contacts. Thus, their entire Cα traces show a RMSD of 0.64 Å (calculated over 169 residues), whereas upon removal of Switch I-interswitch-Switch II residues (39-76), the RMSD decreases to 0.32 Å (calculated over 132 residues).

The two molecules (A and B) in the asymmetric unit of the crystal of GerdArfR1-FL-WT-GDP interact together as a dimer through two equivalent interfaces (Extended Data Fig.6a). Interactions at both interfaces are made between the N-terminal part of the β2-strand of the interswitch (Glu48-Ile63) of one molecule and the N-terminal part of Switch II (Gln69-Arg73) of the second molecule (Extended Data Fig.6b-c). This intermolecular interaction drives Switch II to adopt a β-strand conformation. Altogether, the two interfaces are stabilized by 12 hydrogen bonds as identified using the iterative web server tool PDBePISA^9^. As a consequence, the β2-strand is unzipped from the β3-strand at its base with only a four main-chain hydrogen bond network (Leu54-Leu61) that maintains the tip of the interswitch in a classical position (Extended Data Fig.6d). As a consequence, the unzipping of the β2-β3 sheet displaces the Asp52 (β2-strand) further from the magnesium ion and no longer contributes to its coordination. Thus, in the GerdArfR1-GDP structure, the magnesium ion is shifted by 3.0 Å compared to its classical position in small GTPase structures and exhibits an atypical coordination sphere with Asp65 (β3-strand), Asp24 (P-loop), the Pβ from the GDP, and three water molecules (Extended Data Fig.6e). These differences compared to the classical organization also mean that the Thr29 from the P-loop does not contribute to this atypical nucleotide-binding site coordination sphere. To determine if such a dimer exists in solution, we performed a RALS experiment. However, the results clearly indicate that GerdArfR1-FL-WT-GDP is a monomer in solution (Extended Data Fig.6f). Thus, this structural analysis should be interpreted with caution, as the atypical positions of the Switch I / β2-strand might be a consequence of a crystal packing artifact.

Data collection and refinement statistics for GerdArfR1-Δ12-Q69L-GTP (PDB code 8OUM) and GerdArfR1-FL-WT-GDP (PDB code 8OUN) structures are presented in Extended Data Table 1.

**Extended Data Table 1.** – Crystallographic data collection and refinement statistics.

|  | HodArfR1 |  | GerdArfR1 |  |
| --- | --- | --- | --- | --- |
| Protein |  |  |  |  |
| Nucleotide state | Mg.GTP | Mg.GDP | Mg.GTP | Mg.GDP |
| Fragment, mutant | Δ14, Q72L | FL, WT | Δ12, Q67L | FL, WT |
| PDB code | 8OUK | 8OUL | 8OUM | 8OUN |
| Crystal information |  |  |  |  |
| Space group | *P*4<sub>3</sub>2<sub>1</sub>2 | *P*4<sub>3</sub>2<sub>1</sub>2 | *P*2<sub>1</sub> | *P*2<sub>1</sub> |
| Cell *a*, *b*, *c* (Å) | 59.2, 59.2, 119.7 | 75.7, 75.7, 189.2 | 87.2, 54.9, 121.8 | 43.3, 59.9, 72.3 |
| *α*, *β*, *γ* (°) | 90.0, 90.0, 90.0 | 90.0, 90.0, 90.0 | 90.0, 94.5, 90.0 | 90.0, 102.1, 90.0 |
| No. Mol/ua | 1 | 3 | 5 | 2 |
| Resolution | 1.8 | 3.1 | 2.67 | 1.65 |
| Crystallization Cdt. | 2.0 M Na.Formate HEPES<br>pH 7.5 | 30% PEG4000<br>Tris-HCl pH 8.5 | 8.5% PEG8000<br>Tris pH 7.0 | 18% PEG 8000, Cacodylate<br>pH 6.5 |
|  | - | 0.2M Li2SO4 | 0.1M MgCl2 | 0.2M CaAc |
| Cryo Cdt | 25 % Glycerol | 18% Ethylene Glycol | 25 % Glycerol | 25 % Glycerol |
| Data collection<sup>(a)</sup> |  |  |  |  |
| Synchrotron, Beamline | SOLEIL, PX2 | SOLEIL, PX1 | SOLEIL, PX2 | SOLEIL, PX2 |
| Resolution (Å) | 42.1-1.8 (1.84-1.80) | 70.3-3.10 (3.29-3.10) | 48.0-2.67 (2.83-2.67) | 45.73-1.65 (1.71-1.65) |
| Unique reflections | 20387 (1982) | 10627 (1678) | 32436 (4627) | 39833 (2142) |
| *R*<sub>meas</sub> | 0.044 (0.582) | 0.048 (0.269) | 0.034 (0.260) | 0.070 (0.730) |
| *I* / *σI* | 39.8 (4.4) | 6.2 (1.1) | 6.0 (0.7) | 14.8 (2.1) |
| *CC*<sub>1/2</sub> | 1.000 (0.981) | 0.994 (0.063) | 0.990 (0.446) | 0.999 (0.734) |
| Completeness (%) | 99.0 (97.2) | 99.9 (99.9) | 97.8 (87.1) | 91.8 (99.3) |
| Multiplicity | 25.5 (22.8) | 18.3 (19.1) | 6.8 (5.8) | 5.4 (4.0) |
| Wilson B-factor | 32.9 | 64.7 | 43.6 | 19.6 |
| Refinement |  |  |  |  |
| Resolution (Å) | 42.1-1.8 | 48.5-3.1 | 46.4-2.67 | 45.73-1.65 |
| No. reflections used | 20229 (1947) | 34,801 | 32,148 | 39,822 |
| *R*<sub>work</sub> | 0.186 (0.221) | 0.236 (0.277) | 0.236 (0.383) | 0.206 (0.255) |
| *R*<sub>free</sub> | 0.203 (0.274) | 0.289 (0.389) | 0.267 (0.436) | 0.234 (0.310) |
| *B* factors |  |  |  |  |
| Protein | 42.5 | 61.1 | 68.6 | 23.6 |
| Ligand | 36.0 | 48.4 | 63.9 | 19.4 |
| Solvent | 51.4 | 20.4 | 44.5 | 23.1 |
| Coordinate error (Å) | 0.72 | 0.45 | 0.42 | 0.24 |
| R.m.s.d. |  |  |  |  |
| Bond lengths (Å) | 0.01 | 0.009 | 0.008 | 0.013 |
| Bond angles (°) | 1.62 | 1.00 | 1.0 | 1.63 |
| Ramachandran (%) |  |  |  |  |
| Favored region | 96.5 | 91.9 | 95.6 | 96.6 |
| Allowed region | 3.5 | 7.4 | 4.2 | 3.4 |
| Outliers | 0 | 0.7 | 0.2 | 0 |
| Number of atoms |  |  |  |  |
| Protein | 1377 | 4139 | 6450 | 2878 |
| Ligand | 33 | 87 | 165 | 2 |
| Solvent | 97 | 9 | 122 | 313 |

<sup>(a)</sup>Data was collected on a single crystal. Values in parentheses are for the highest-resolution shell.

**Extended Data Table 2.** – Crystallographic information from crystal structures of the GDP-bound form of Arf proteins used for structural analysis.

|  | PDB code | Res.(Å) | SG | Unit-cell parameters |  |  |  | Mol/ua | ref |
| --- | --- | --- | --- | --- | --- | --- | --- | --- | --- |
| | | | | *a* (Å) | *b*(Å) | *c* (Å) | $\beta$ (°) | | |
| **HodArfR1** | 8OUL | 3.1 | *P*4<sub>3</sub>2<sub>1</sub>2 | 75.7 | 75.7 | 189.3 | 90.0 | 3 | This study |
| **GerdArfR1** | 8OUN | 1.7 | *P*2<sub>1</sub> | 43.3 | 59.9 | 72.3 | 102.1 | 2 | This study |
| **HsArf1** | 1HUR | 2.0 | *C*2 | 121.0 | 45.9 | 91.8 | 133.5 | 2 | Amor et al., 1994<sup>10</sup> |
| **HsArl8** | 4ILE | 2.7 | *P*2<sub>1</sub>2<sub>1</sub>2 | 118.0 | 37.4 | 39.3 | 90.0 | 1 | No publication |
| **LmArl1** | 2X77 | 2.1 | *P*2<sub>1</sub> | 35.3 | 78.9 | 58.0 | 93.1 | 2 | Fleming et al., 2010<sup>11</sup> |

**Extended Data Table 3.** Plasmids for yeast expression used in this study.

| Plasmid | Vector-Promoter-Expressed protein | Reference |
| --- | --- | --- |
| pCLJ1101 | pRS416-prADH-GFP | S. Leon |
| pCLJ1103 | pRS416-prADH-HodArfR1-GFP | This study |
| pCLJ1104 | pRS416-prADH-HodArfR1-S31N-GFP | This study |
| pCLJ1105 | pRS416-prADH-HodArfR1-Q72L-GFP | This study |
| pCLJ1111 | pRS416-prADH-HodArfR1-ΔN(1-15)-GFP | This study |
| pCLJ1112 | pRS416-prADH-HodArfR1-Q72L-ΔN(1-15)-GFP | This study |
| pCLJ1117 | pRS416-prADH-GerdArfR1-GFP | This study |
| pCLJ1118 | pRS416-prADH-GerdArfR1-T27N-GFP | This study |
| pCLJ1119 | pRS416-prADH-GerdArfR1-Q69L-GFP | This study |
| pCLJ1120 | pRS416-prADH-GerdArfR1-ΔN(1-15)-GFP | This study |
| pHCMM4 | pCYC1-prCYC1-Sec61-mCherry | Medina-Munoz et al., 2020<sup>12</sup> |

**Extended Data Figure 1.**
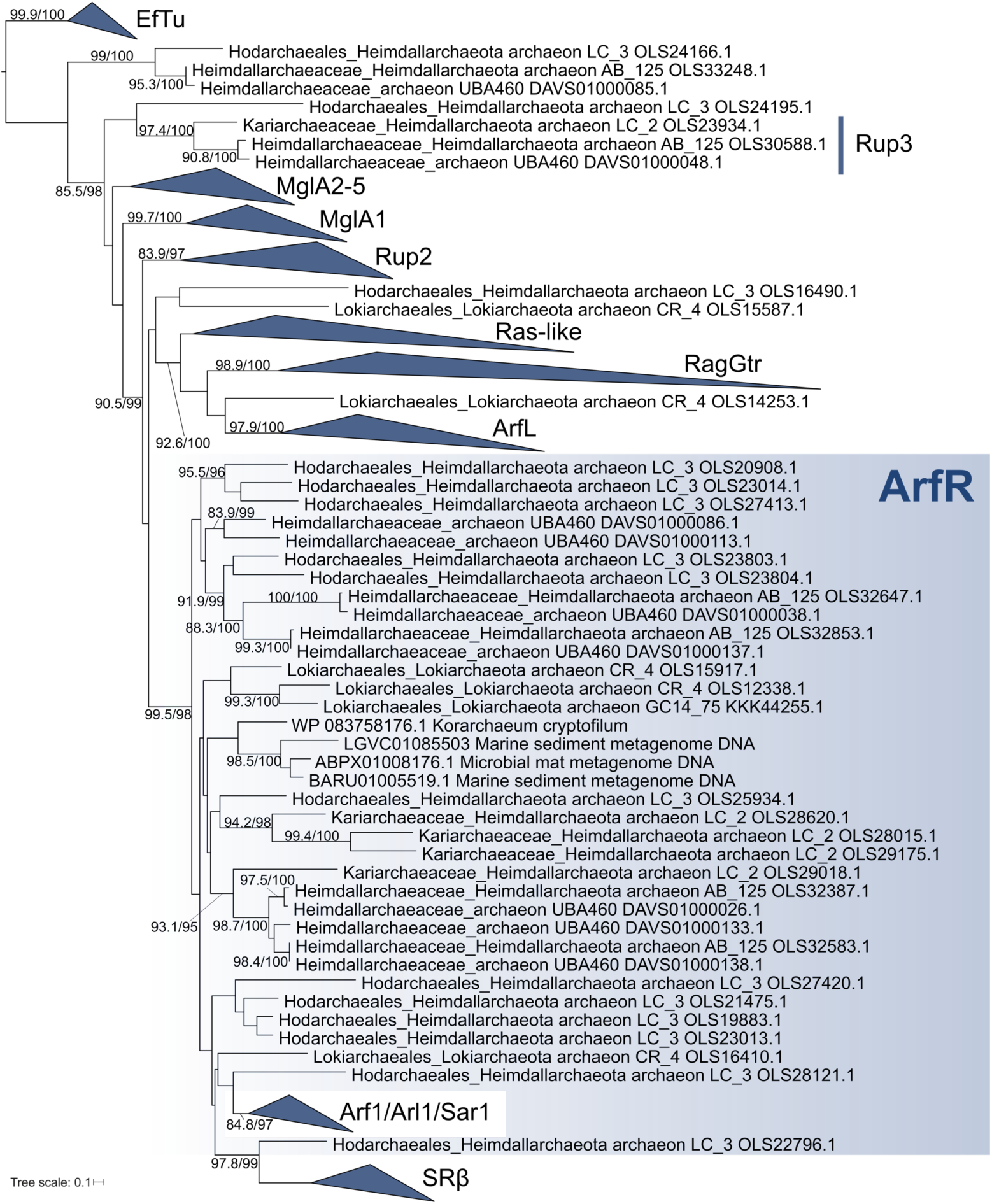
– Initial phylogenetic analysis that pointed to the existence of ArfR, a group of asgardarchaeote GTPases specifically related to eukaryotic Arf family and SRβ. The tree was inferred using IQ-TREE with the LG+F+I+G4 model. Numbers at branches (shown when >80/95) correspond to SH-aLRT support values (right side of the slash) and ultrafast bootstrap support values (left side of the slash) calculated from 10000 replicates. As the basis for this analysis we used the sequence set utilized previously by Klinger et al. (2016)^13^, which included the Ras superfamily GTPase set encoded by a single asgardarchaeote MAG (the very first reported, i.e. “Lokiarchaeota archaeon GC14_75”). The dataset was expanded by addition of Ras superfamily sequences from new genome assemblies from the Asgard superphylum^14,15^, the korarchaeian *Candidatus* Korarchaeum cryptofilum, and three related sequences of an unknown provenance identified in metagenome assemblies at the early stage of the project (most likely corresponding to fragments derived from Asgardarchaeota genomes). EF-Tu sequences (GTPases not belonging to the Ras superfamily) were considered as an outgroup. For the sake of simplicity, GTPase clades observed and defined in the previous study^13^ were collapsed as triangles. The clade denoted “Ras-like” contains eukaryotic Rab/Ras/Rho/Ran sequences, prokaryotic Rup1 sequences, and asgardarchaeote RasL as defined by Klinger et al. (2016)^13^. The novel group denoted “ArfR” (Arf-related) consists of archaebacterial GTPases specifically related to eukaryotic Arf, Arl, Sar1 and SRβ proteins, whereas Rup3 denotes another novel GTPase group, which is specific for Heimdallarchaeia (see Fig. S2 for a more detailed analysis). Note that the existence of ArfR was missed in Klinger et al. (2016), because the single ArfR protein encoded by Lokiarchaeota archaeon GC14_75 (Lokiarch_19980; GenBank KKK44255.1) was removed from analysis for being considered too divergent.

**Extended Data Figure 2.**
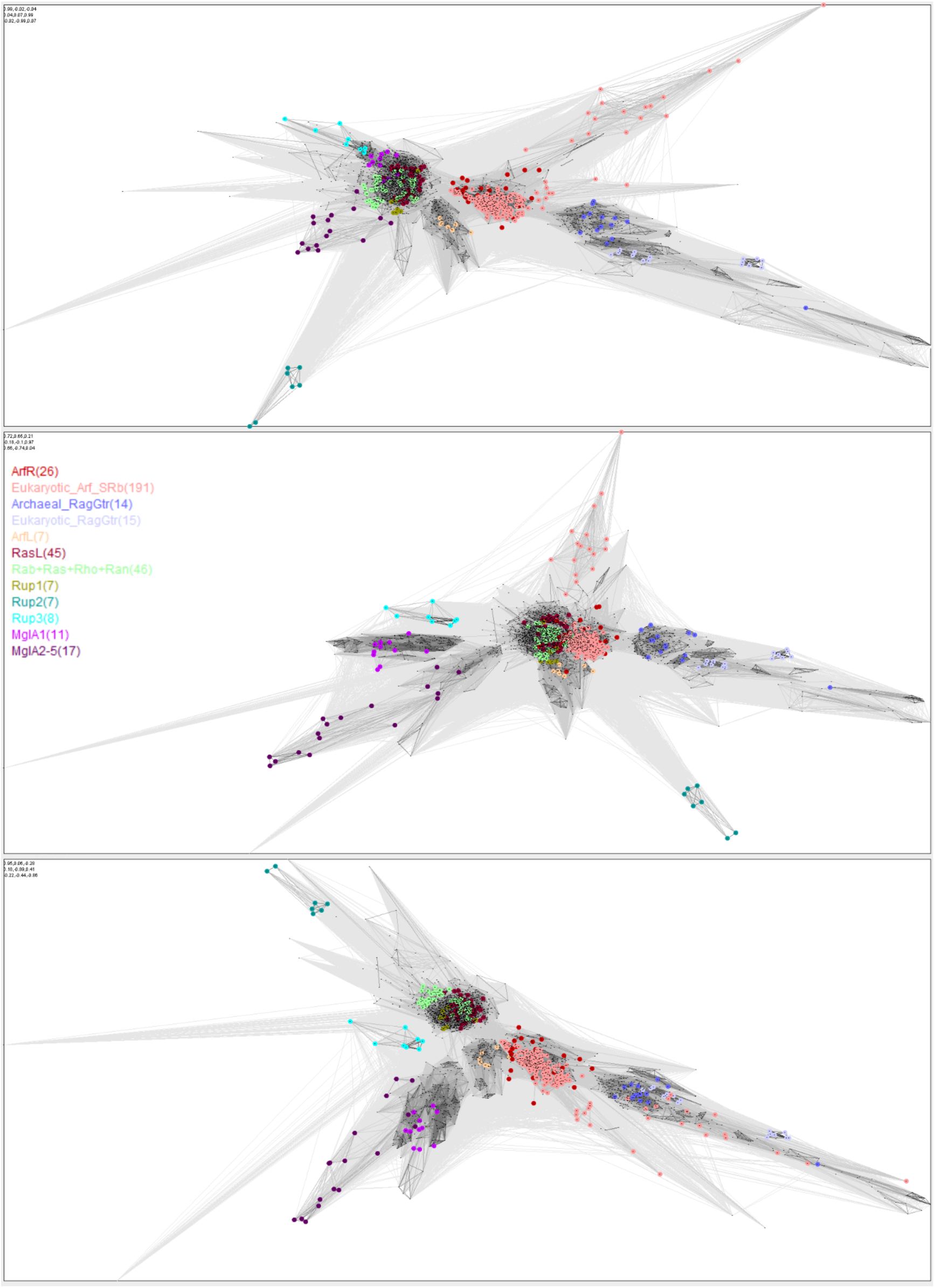
– Sequence diversity in the Ras superfamily analysed with CLANS. Displayed are three different 2D projections of a 3D plot showing the results of clustering of 2994 Ras superfamily protein sequences (trimmed to the GTPase domain) based on their sequence similarity scored by all-to-all blastp comparisons. Individual sequences are represented by vertices (dots) connected by edges when their high scoring segment pairs value is ≤ 10^-5^. Reference sequences from different predefined subgroups are highlighted with a subgroup-specific colour (according to the graphical legend provided in the middle panel). Note the clear separation from other GTPase subgroups of a cluster comprised of archaeal ArfR sequences and eukaryotic Arf family and SRβ sequences, with some of the more divergent eukaryotic sequences (e.g. Arl16 or SRβ) situated further away from the core of the cluster towards the periphery. Similarly, the cluster comprised of archaeal (asgradarchaeote) and eukaryotic RagGtr sequences includes some more divergent members connecting to the core from a distance. Note the tight packing of the eukaryotic Rab/Ras/Rho/Ran sequences and the prokaryotic Rup1 group with the asgardarchaeote RasL sequences and the separate status of the newly defined Rup3 group.

**Extended Data Figure 3.**
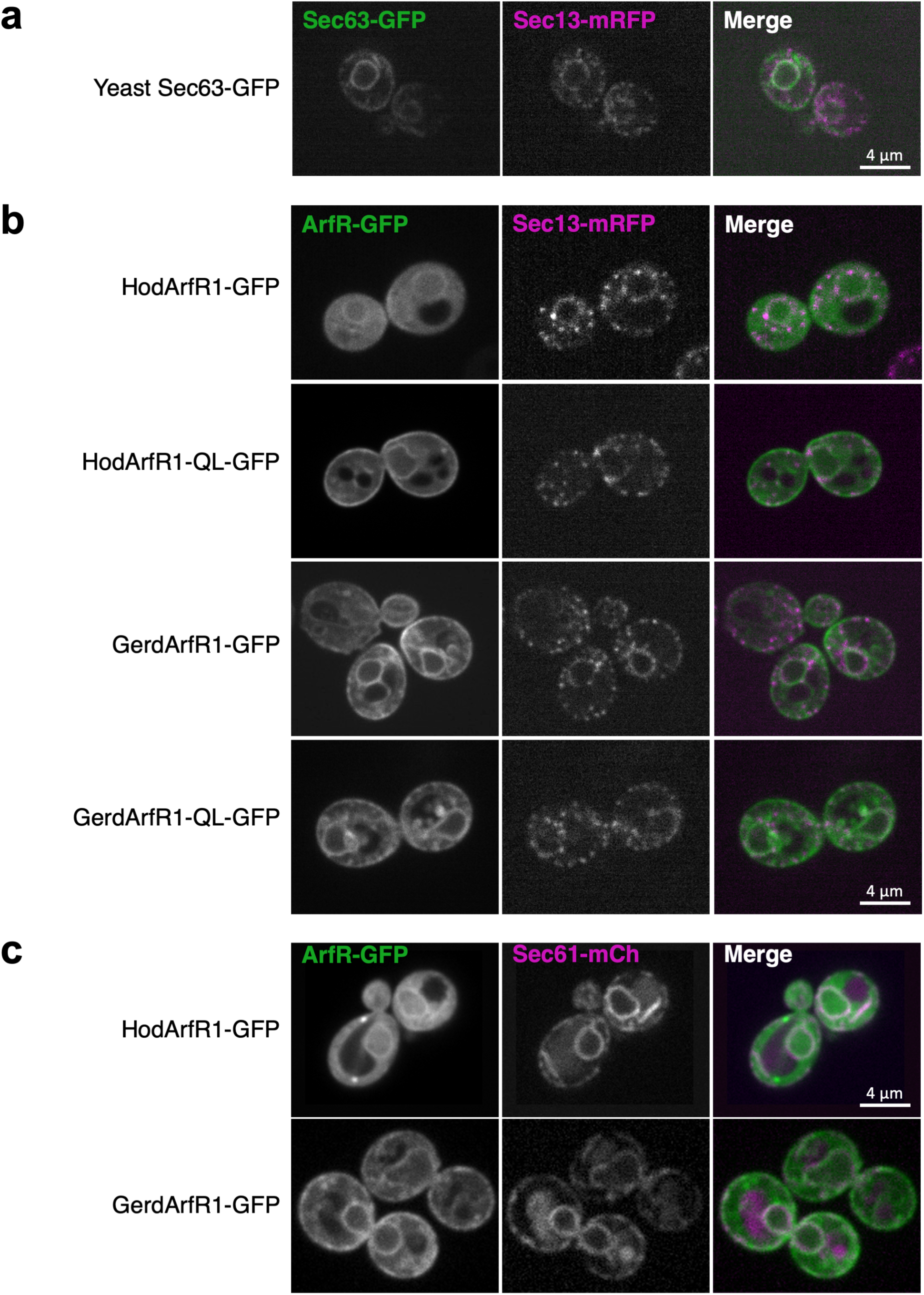
– Asgardarchaeote ArfR proteins localize to the ER in yeast cells. (**a**) Sec13-mRFP localizes to the ER and ER exit sites. The ER protein Sec63-GFP, a component of the ER protein translocation channel, was expressed in in the yeast strain EY0987 *Sec13::mRFP MATα his3Δ1 leu2Δ0 lys2Δ0 ura3Δ0 Sec13::mRFP-kanMX6*, in which Sec13 is fused to mRFP at its C-terminus in its chromosomal locus^16^. Sec63-GFP (left panel), the COPII subunit Sec13-mRFP (centre panel) and the merged image (left panel) are shown. Sec13-mRFP labels both the ER and ER exit sites, which are localized as discrete puncta along the ER membrane. **(b)** HodArfR1 and GerdArfR1 localize to the ER, marked by Sec13-mRFP. Wild type HodArfR1, HodArfR1-Q72L, wild type GerdArfR1 and GerdArfR1-Q69L were expressed as GFP fusion proteins in the yeast strain EY0987 *Sec13::mRFP MATα his3Δ1 leu2Δ0 lys2Δ0 ura3Δ0 Sec13::mRFP-kanMX6*. Representative images of the indicated GFP-tagged ArfR proteins (left panels) and the same cells expressing Sec13-mRFP (centre panels) are shown. The merge image is shown in the right panels. **(c)** HodArfR1 and GerdArfR1 localize to the ER, marked by Sec61-mCherry, the ER protein translocation channel. Wild type HodArfR1 and wild type GerdArfR1 were expressed as GFP fusion proteins in the yeast strain BY4742 *MATα his3Δ1 leu2Δ0 lys2Δ0 ura3Δ0*, which was co-transformed with pHCMM4 Sec61- mCherry. Representative images of the indicated GFP-tagged ArfR proteins (left panels) and the same cells expressing Sec61-mCherry (centre panels) are shown. The merge image is shown in the right panels.

**Extended Data Figure 4.**
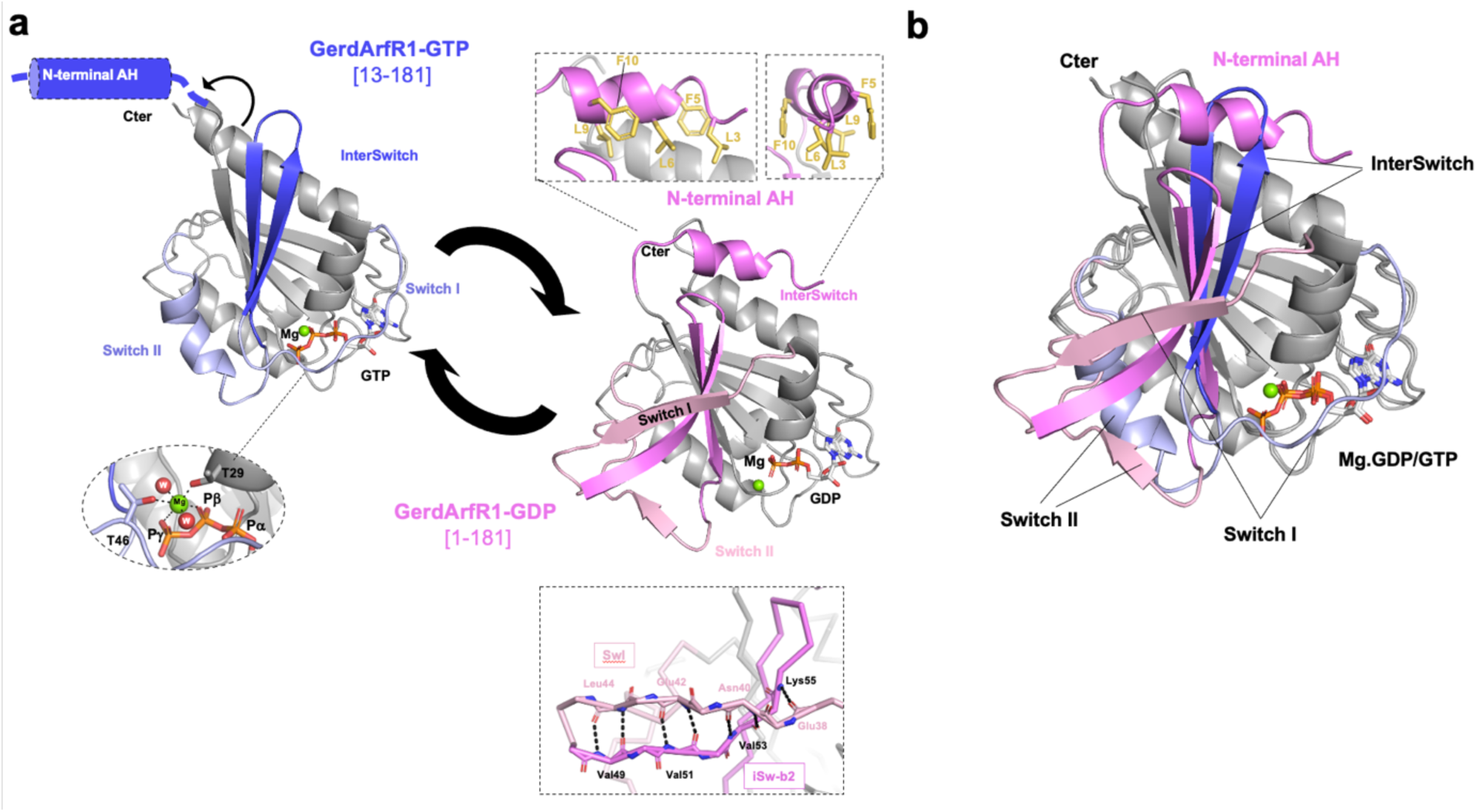
– GDP/GTP structural cycle of GerdArfR1. (**a**) (left) Crystal structure of GerArfR1-Δ12-Q69L bound to Mg.GTP. The protein is shown with the interswitch in blue and Switch I/Switch II in light blue. Below, the nucleotide binding site of the Mg.GTP is enlarged. The N-terminal helix, which is absent from the fragment crystallized, is shown schematically. (right) Crystal structure of GerdArfR1-FL-WT bound to Mg.GDP. The protein is shown with the N-terminal helix and the interswitch in pink and Switch I/Switch II in light pink. Above, the N-terminal helix is magnified and shown in two orientations rotated by 90°. Hydrophobic residues Leu3, Phe5, Leu6, Leu9 and Phe10 from the N-terminal AH of GerdArfR1 (indicated in yellow) are buried in the hydrophobic pocket formed by the tip of the interswitch and the C-terminal helix. Overall, the protein is shown with a cartoon representation, the magnesium is indicated by a green sphere and the GTP is shown with a stick representation. **(b)** Superimposition of the GTP-bound and GDP-bound forms of HodArfR1. Colors are as described in part (a).

**Extended Data Figure 5.**
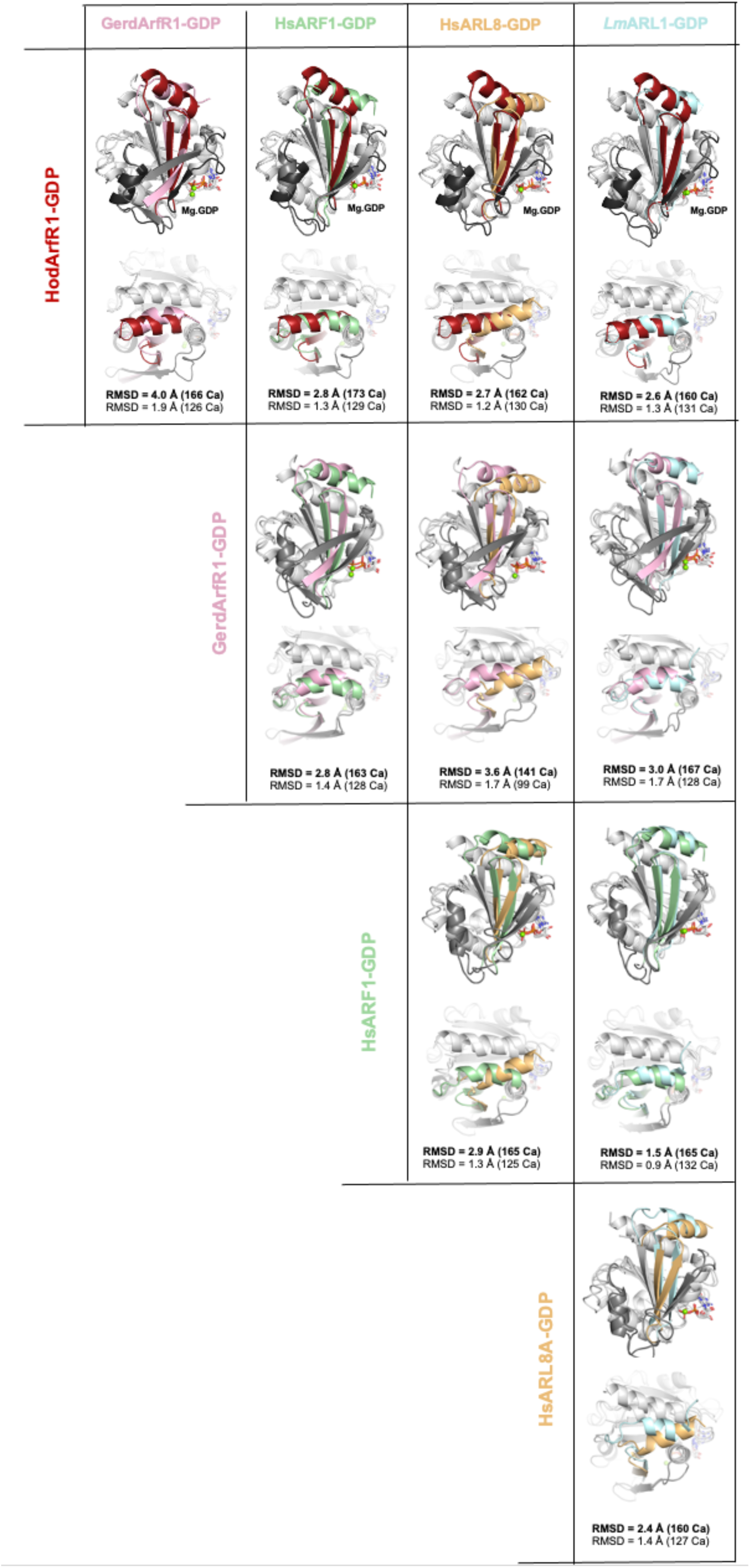
– Structural Tables for the comparison of the GDP-bound forms of asgardarchaeote and eukaryotic Arf proteins. HodArfR1-GDP, GerdArfR1-GDP, HsArf1-GDP, HsArl8-GDP and LmArl1-GDP are compared to each other in pairs. **(a)** ‘face” orientation showing the N-terminal and the interswitch regions in colors and Switch I/Switch II regions in grey and below, **(b)** a 90° “top” rotation focusing on the N-terminal helix. Superimposition was performed on residues of the P-loop, strands β1-β3 and the C-terminal helix. The same orientation is shown for all superimpositions. Hod, Hodarchaeales; Hs, *Homo sapiens*; Lm, *Leishmania major*.

**Extended Data Figure 6.**
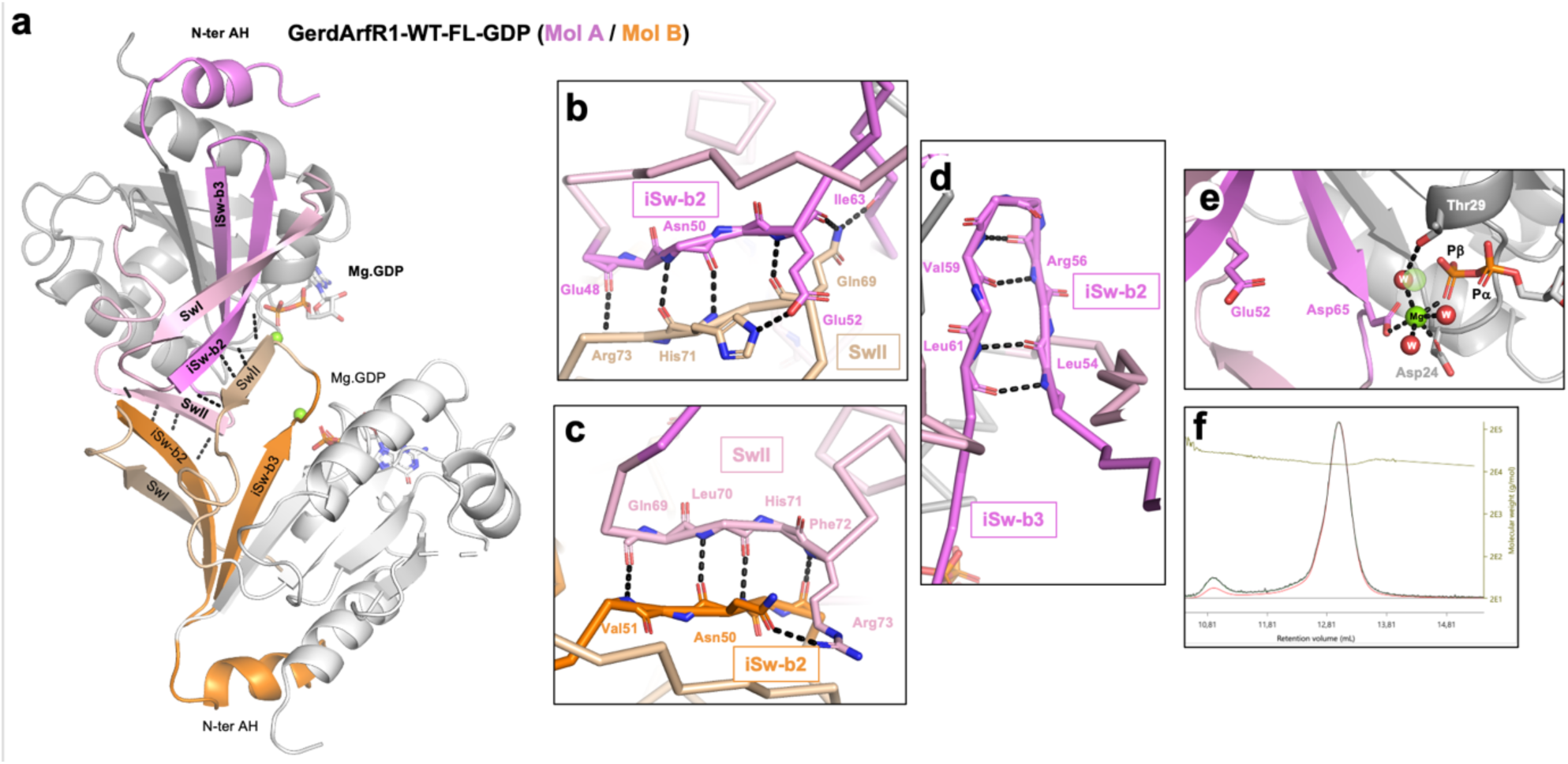
Crystal dimer of GerdArfR1-GDP. (**a**) Overall organization of the crystal dimer of GerdArfR1-GDP. The N-terminal AH and switch regions of molecule A are in pink and those of molecule B are in orange. **(b-c)** Detailed views of the two interfaces of the dimer involving Switch II of one molecule and the β2-strand of the interswitch from the other molecule. **(d)** The hydrogen network of the interswitch (β2-β3 strands) shows an unzipping compared to other Arf-GDP structures. **(e)** The atypical coordination sphere of the magnesium ion. The classical position of the magnesium ion is indicated by a transparent green sphere. **(f)** Sec-RALS experiment showing that in solution GerdArfR1-GDP is monomeric.

