## Supplementary Figures for "Arf family GTPases are present in Asgard archaea"

**Fig. S1. The phylogenetic position of the Asgardarchaeota genomes selected for analysis.** The tree was inferred using FastTree (with default settings) from a multiple alignment of sequences of the DNA-dependent RNA polymerase subunit B (RPOB) used as a phylogenetic marker. The major asgardarchaeote groups are highlighted, using the original nomenclature considering Asgard archaea as a superphylum consisting of multiple phyla. Note that Njordarchaeota (presently classified as the order Njordarchaeales in the class Heimdallarchaeia) do not branch together with the other asgardarchaeote lineages in the tree but appear nested in the TACK group (phylum Thermoproteota) as a sister clade of Korarchaeota (=class Koraracheaia). This seems to reflect a previously detected artificial phylogenetic signal attracting Njordarchaeales to Korarchaeia due to a convergently shared sequence composition bias (Eme et al. 2023)[9]. The genomes, for which Ras superfamily GTPases were identified and analysed in this study (Table S2), are highlighted, except for a few MAGs that lack a RPOB gene (obviously due to their incompleteness; see Methods for further details).

**Fig. S2. Details on Rup3, a novel Heimdallarchaeia-specific GTPase group.** This group joins the two previously defined Rup groups ("Rup" standing for Ras superfamily GTPase of unknown function in prokaryotes) defined by Wuichet and Søgaard-Andersenet (2014)[42]. (a) The tree was inferred using IQ-TREE (with the LG+I+G4 model) from the same set of archaeal Ras superfamily sequences as used for obtaining the tree displayed as Fig. S4 but expanded by the addition of the sequences identified here to belong to the Rup3 group. Note that for this analysis we searched for the occurrence of Rup3 genes beyond the 62 reference asgardarchaeote genome assemblies selected for the other analyses reported in this paper, but identified them only in assemblies representing different lineages of Heimdallarchaeia. The tree confirms the monophyly of the Rup3 group and its separation from the other major groups of Ras superfamily GTPases. Branch support was assessed by the SH-aLRT test and the ultrafast bootstrap algorithm (both 1000 replicates). (b) A segment of a multiple alignment including Rup3 sequences (bottom) together with sequences representative of other major Ras superfamily groups. Conserved motifs of the GTPase domain, i.e. the boxes G1 (P-loop), G2 (a conserved threonine residue), and G3 (Walker B motif) are shown according to their position in the human Arf1 sequence as detected by Conserved Domain Database search at NCBI. Note the sequence expansions between the G1 and G2 boxes and between the G2 and G3 boxes are unique for Rup3 sequences.

**Fig. S3. Phylogenetic analysis of Ras superfamily GTPases from Asgardarchaeota.** The tree was inferred from a multiple alignment of 2626 sequences using IQ-TREE 1.5.5 with the LG+F+R10 substitution model. Branch support was assessed by the SH-aLRT test with 1000 replicates. Sequences from the very first asgardarchaeote MAG reported ("Lokiarchaeota archaeon GC14_75") are marked with a green star or a red cross depending on whether they were included or excluded, respectively, from the previous phylogenetic analysis by Klinger et al. (2016)[22].

**Fig. S4. Full version of the tree resulting from a phylogenetic analysis of a representative dataset of the Ras superfamily sequences from archaea and reference eukaryotic and eubacterial sequences.** The tree was inferred using IQ-TREE with the LG+I+G4 model from a multiple alignment of 996 sequences. The numbers at branches correspond to support values calculated with the SH-aLRT test and the ultrafast bootstrap algorithm (both 1000 replicates). Major GTPase subgroups are annotated in the tree, including the different paralogs of the eukaryotic family as defined previously (Vargová et al. 2021)[23]. Note that the monophyly of the eukaryotic Arf family (including SRβ) is disrupted by one of the asgardarchaeote ArfR sequences (from a Hodarchaeales member) nested among them, branching specifically with SRβ. Another hodarchaealean sequence (OLS22796.1; not included in the tree displayed in the figure) branches with SRβ in the tree shown in Extended Data Figure 1. While this may suggest that SRβ evolved from a separate (hodarchaealean) ArfR gene than the (other) Arf family members, the specific relationships between the eukaryotic Arf/SRβ and the archaeal ArfR proteins turned out to be unstable in additional phylogenetic analyses focusing on this question.

**Fig. S5. Occurrence of ArfR in Korarchaeia.** The tree shows phylogenetic relationships among korarchaeial genomes as inferred from RPOB protein sequences encoded by them. The tree was inferred using IQ-TREE 1.5.5, with LG+F+R7 selected as the best-fit substitution model. Numbers at branches correspond to SH-aLRT support values calculated from 1000 replicates. The accession numbers indicated in the tip labels correspond to the scaffolds (contigs) in the respective genome assemblies or MAGs that contain the RPOB gene. In a few cases two separate genome sequence fragments were joined (indicated by the "+" sign) to obtain a complete (or longer) RPOB gene sequence. Genomes in which ArfR genes were identified are highlighted in red and the branch where an ArfR gene was acquired by horizontal transfer from an asgardarchaeote is marked with an arrow.

**Fig. S6. Phylogenetic relationship between korarchaeial and asgardarchaeote ArfR proteins.** Displayed is a simplified version of a tree (with the non-focal GTPase groups collapsed as triangles) inferred using IQ-TREE (with the substitution model LG+I+G4) from the dataset used to infer the tree displayed in Fig. S4 but expanded to include all ArfR sequences identified in presently available korarchaeial genome assemblies. All korarchaeial sequences form a monophyletic group nested among asgardarchaeote ArfR sequences. Note that some korarchaeial genomes contain two different ArfR paralogs, as evident from the sequence IDs – for instance, JALUEI010000145.1 and JALUEI010000136.1 are ArfR genes at two different scaffolds in the assembly JALUEI000000000.1.

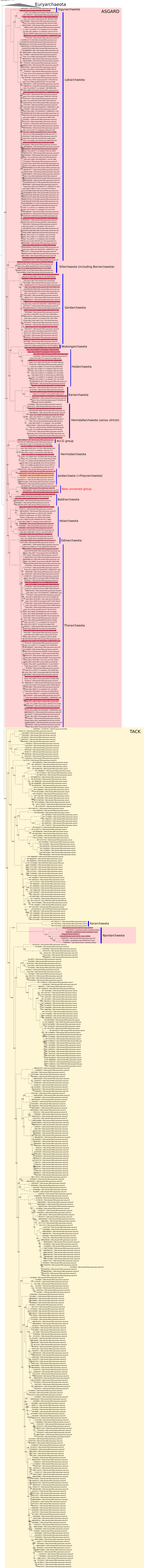
Euryarchaeota
ASGARD
Gigunarchaeota
Lokiarchaeota
Sifarchaeota (including Borrarchaeota)
Gerdarchaeota
Wukongarchaeota
Hodarchaeota
Kariarchaeota
Heimdallarchaeota (sensu stricto)
ALCG group
Hermodarchaeota
Jordarchaeia (=Freyarchaeota)
New uncurated group
Baldrarchaeota
Helarchaeota
Odinarchaeota
Thorarchaeota
TACK
Korarchaeota
Njordarchaeota

A)

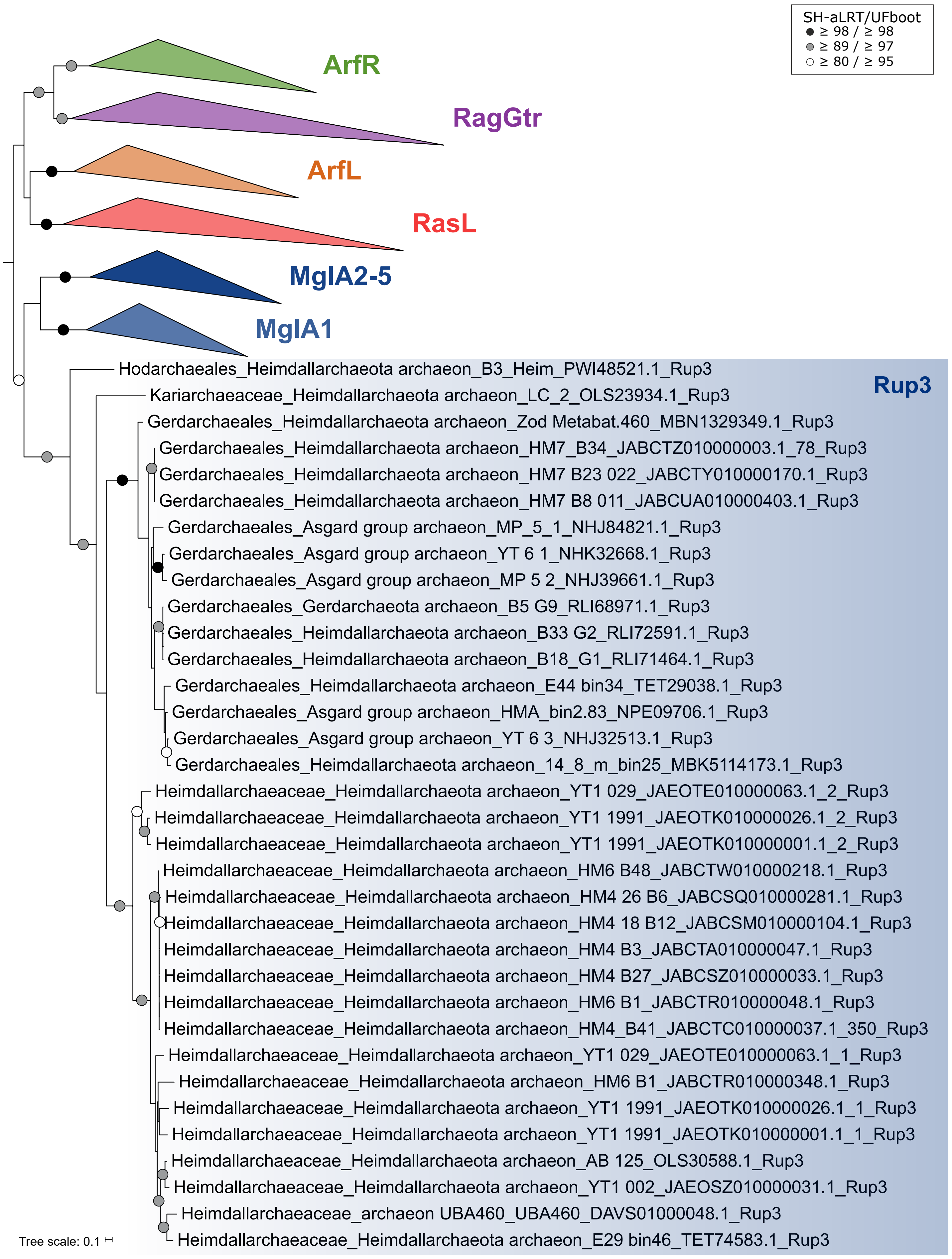
SH-aLRT/UFboot
≥ 98 / ≥ 98
≥ 89 / ≥ 97
≥ 80 / ≥ 95
ArfR
RagGtr
ArfL
RasL
MglA2-5
MglA1
Rup3
Hodarchaeales_Heimdallarchaeota archaeon_B3_Heim_PWI48521.1_Rup3
Kariarchaeaceae_Heimdallarchaeota archaeon_LC_2_OLS23934.1_Rup3
Gerdarchaeales_Heimdallarchaeota archaeon_Zod Metabat.460_MBN1329349.1_Rup3
Gerdarchaeales_Heimdallarchaeota archaeon_HM7_B34_JABCTZ010000003.1_78_Rup3
Gerdarchaeales_Heimdallarchaeota archaeon_HM7 B23 022_JABCTY010000170.1_Rup3
Gerdarchaeales_Heimdallarchaeota archaeon_HM7 B8 011_JABCUA010000403.1_Rup3
Gerdarchaeales_Asgard group archaeon_MP_5_1_NHJ84821.1_Rup3
Gerdarchaeales_Asgard group archaeon_YT 6 1_NHK32668.1_Rup3
Gerdarchaeales_Asgard group archaeon_MP 5 2_NHJ39661.1_Rup3
Gerdarchaeales_Gerdarchaeota archaeon_B5 G9_RLI68971.1_Rup3
Gerdarchaeales_Heimdallarchaeota archaeon_B33 G2_RLI72591.1_Rup3
Gerdarchaeales_Heimdallarchaeota archaeon_B18_G1_RLI71464.1_Rup3
Gerdarchaeales_Heimdallarchaeota archaeon_E44 bin34_TET29038.1_Rup3
Gerdarchaeales_Asgard group archaeon_HMA_bin2.83_NPE09706.1_Rup3
Gerdarchaeales_Asgard group archaeon_YT 6 3_NHJ32513.1_Rup3
Gerdarchaeales_Heimdallarchaeota archaeon_14_8_m_bin25_MBK5114173.1_Rup3
Heimdallarchaeaceae_Heimdallarchaeota archaeon_YT1 029_JAEOTE010000063.1_2_Rup3
Heimdallarchaeaceae_Heimdallarchaeota archaeon_YT1 1991_JAEOTK010000026.1_2_Rup3
Heimdallarchaeaceae_Heimdallarchaeota archaeon_YT1 1991_JAEOTK010000001.1_2_Rup3
Heimdallarchaeaceae_Heimdallarchaeota archaeon_HM6 B48_JABCTW010000218.1_Rup3
Heimdallarchaeaceae_Heimdallarchaeota archaeon_HM4 26 B6_JABCSQ010000281.1_Rup3
Heimdallarchaeaceae_Heimdallarchaeota archaeon_HM4 18 B12_JABCSM010000104.1_Rup3
Heimdallarchaeaceae_Heimdallarchaeota archaeon_HM4 B3_JABCTA010000047.1_Rup3
Heimdallarchaeaceae_Heimdallarchaeota archaeon_HM4 B27_JABCSZ010000033.1_Rup3
Heimdallarchaeaceae_Heimdallarchaeota archaeon_HM6 B1_JABCTR010000048.1_Rup3
Heimdallarchaeaceae_Heimdallarchaeota archaeon_HM4_B41_JABCTC010000037.1_350_Rup3
Heimdallarchaeaceae_Heimdallarchaeota archaeon_YT1 029_JAEOTE010000063.1_1_Rup3
Heimdallarchaeaceae_Heimdallarchaeota archaeon_HM6 B1_JABCTR010000348.1_Rup3
Heimdallarchaeaceae_Heimdallarchaeota archaeon_YT1 1991_JAEOTK010000026.1_1_Rup3
Heimdallarchaeaceae_Heimdallarchaeota archaeon_YT1 1991_JAEOTK010000001.1_1_Rup3
Heimdallarchaeaceae_Heimdallarchaeota archaeon_AB 125_OLS30588.1_Rup3
Heimdallarchaeaceae_Heimdallarchaeota archaeon_YT1 002_JAEOSZ010000031.1_Rup3
Heimdallarchaeaceae_archaeon UBA460_UBA460_DAVS01000048.1_Rup3
Heimdallarchaeaceae_Heimdallarchaeota archaeon_E29 bin46_TET74583.1_Rup3
Tree scale: 0.1

B)

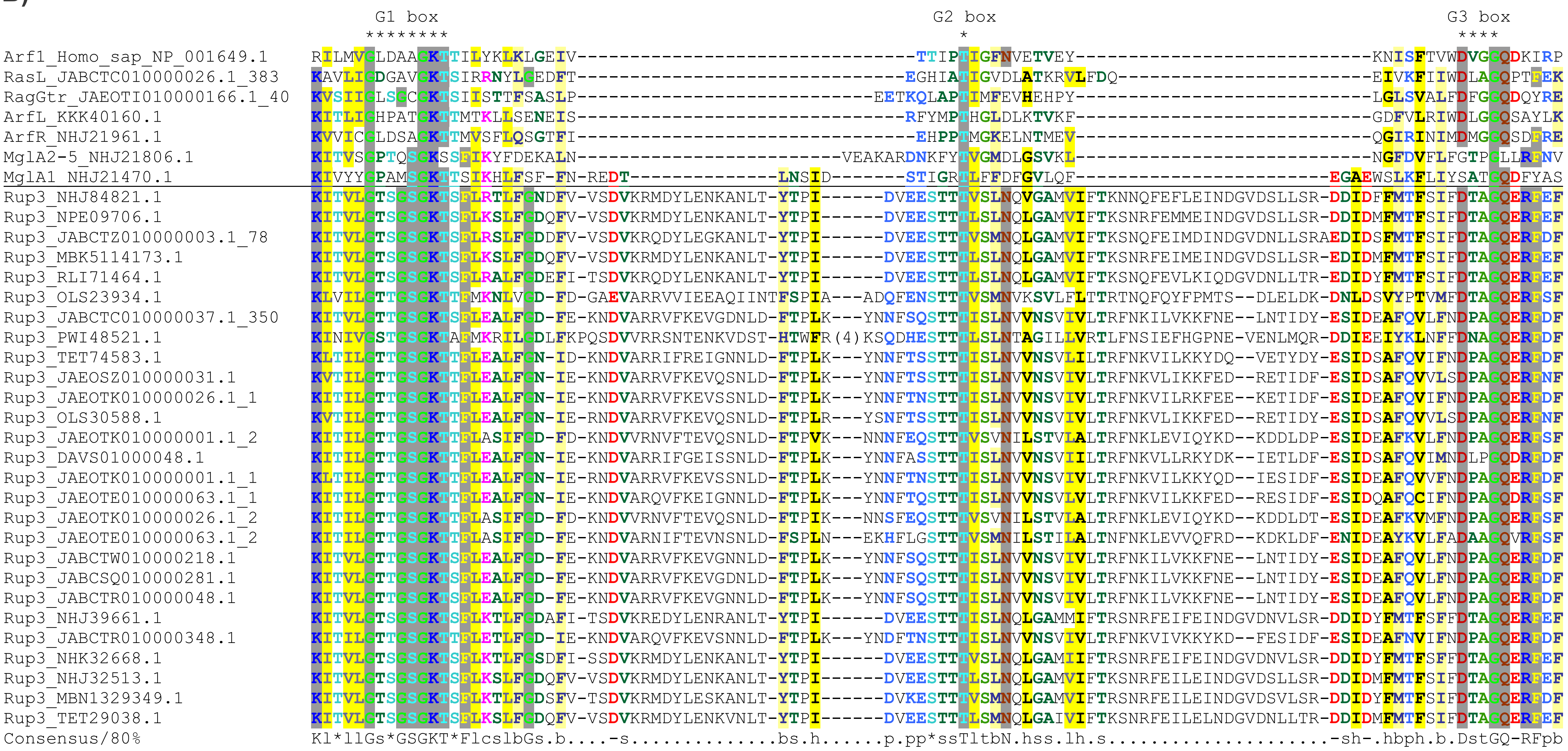
G1 box
G2 box
G3 box
Arf1_Homo_sap_NP_001649.1
RasL_JABCTC010000026.1_383
RagGtr_JAEOTI010000166.1_40
ArfL_KKK40160.1
ArfR_NHJ21961.1
MglA2-5_NHJ21806.1
MglA1_NHJ21470.1
Rup3_NHJ84821.1
Rup3_NPE09706.1
Rup3_JABCTZ010000003.1_78
Rup3_MBK5114173.1
Rup3_RLI71464.1
Rup3_OLS23934.1
Rup3_JABCTC010000037.1_350
Rup3_PWI48521.1
Rup3_TET74583.1
Rup3_JAEOSZ010000031.1
Rup3_JAEOTK010000026.1_1
Rup3_OLS30588.1
Rup3_JAEOTK010000001.1_2
Rup3_DAVS01000048.1
Rup3_JAEOTK010000001.1_1
Rup3_JAEOTE010000063.1_1
Rup3_JAEOTK010000026.1_2
Rup3_JAEOTE010000063.1_2
Rup3_JABCTW010000218.1
Rup3_JABCSQ010000281.1
Rup3_JABCTR010000048.1
Rup3_NHJ39661.1
Rup3_JABCTR010000348.1
Rup3_NHK32668.1
Rup3_NHJ32513.1
Rup3_MBN1329349.1
Rup3_TET29038.1
Consensus/80%

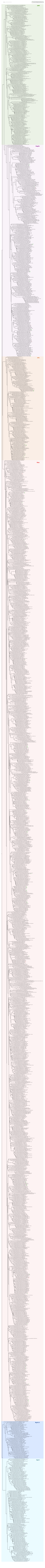
Tree scale: 1
included in the final phylogenetic analysis in Klinger et al. 2016
not included in the final phylogenetic analysis in Klinger et al. 2016
ArfR
RagGtr
ArfL
RasL
MglA2-5
MglA1

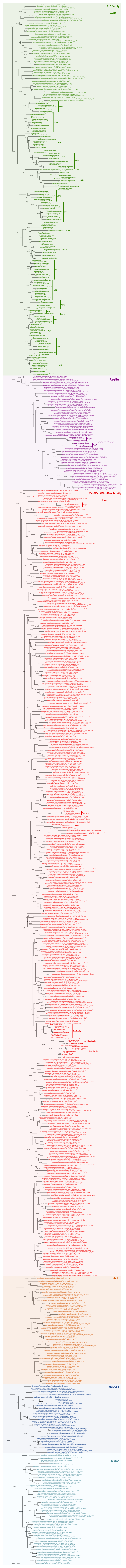
Arf family
+
ArfR
RagGtr
Rab/Ran/Rho/Ras family
+
RasL
ArfL
MglA2-5
MglA1

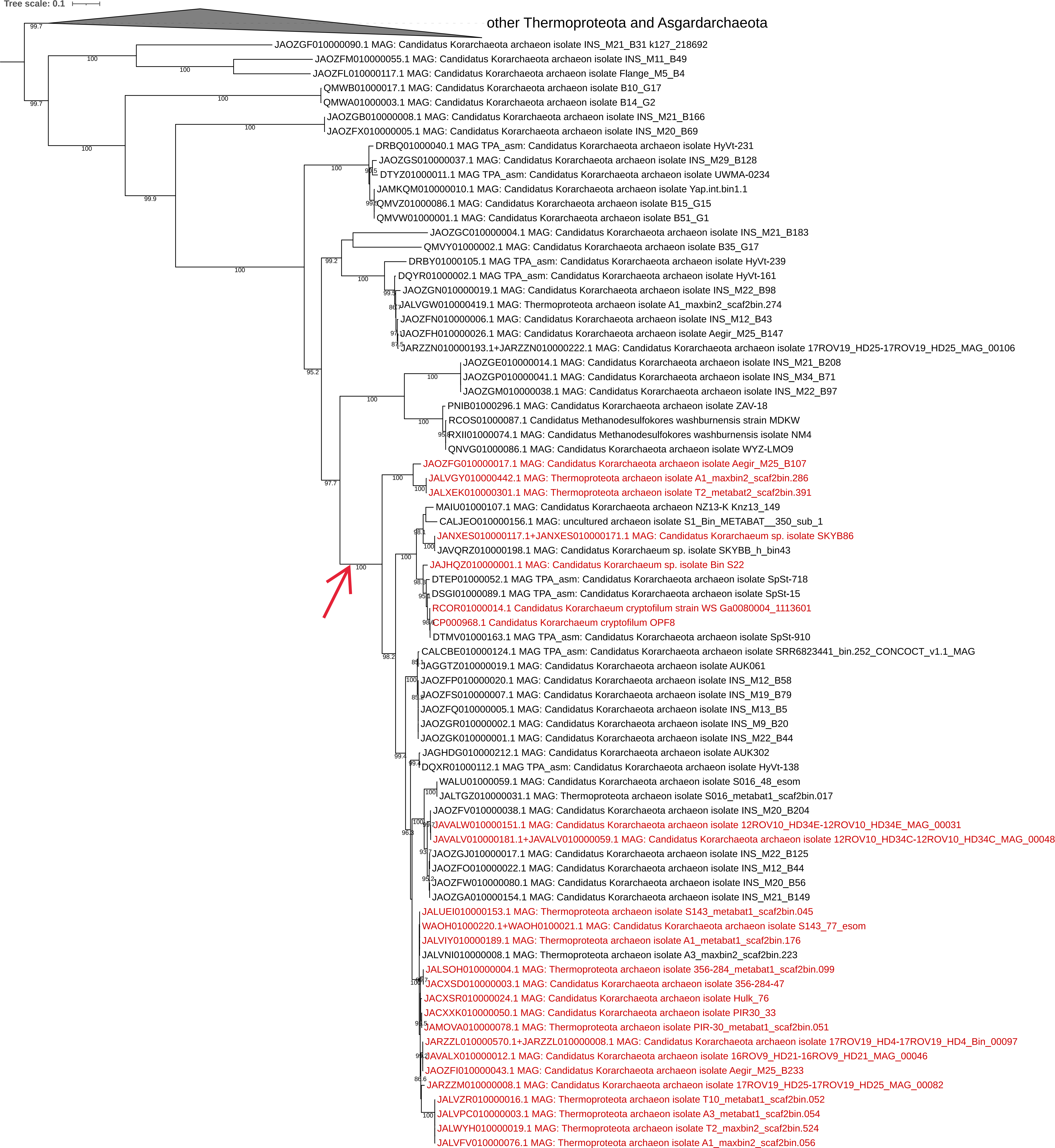

Tree scale: 0.1
other Thermoproteota and Asgardarchaeota
JAOZGF010000090.1 MAG: Candidatus Korarchaeota archaeon isolate INS_M21_B31 k127_218692
JAOZFM010000055.1 MAG: Candidatus Korarchaeota archaeon isolate INS_M11_B49
JAOZFL010000117.1 MAG: Candidatus Korarchaeota archaeon isolate Flange_M5_B4
QMWB01000017.1 MAG: Candidatus Korarchaeota archaeon isolate B10_G17
QMWA01000003.1 MAG: Candidatus Korarchaeota archaeon isolate B14_G2
JAOZGB010000008.1 MAG: Candidatus Korarchaeota archaeon isolate INS_M21_B166
JAOZFX010000005.1 MAG: Candidatus Korarchaeota archaeon isolate INS_M20_B69
DRBQ01000040.1 MAG TPA_asm: Candidatus Korarchaeota archaeon isolate HyVt-231
JAOZGS010000037.1 MAG: Candidatus Korarchaeota archaeon isolate INS_M29_B128
DTYZ01000011.1 MAG TPA_asm: Candidatus Korarchaeota archaeon isolate UWMA-0234
JAMKQM010000010.1 MAG: Candidatus Korarchaeota archaeon isolate Yap.int.bin1.1
QMVZ01000086.1 MAG: Candidatus Korarchaeota archaeon isolate B15_G15
QMVW01000001.1 MAG: Candidatus Korarchaeota archaeon isolate B51_G1
JAOZGC010000004.1 MAG: Candidatus Korarchaeota archaeon isolate INS_M21_B183
QMVY01000002.1 MAG: Candidatus Korarchaeota archaeon isolate B35_G17
DRBY01000105.1 MAG TPA_asm: Candidatus Korarchaeota archaeon isolate HyVt-239
DQYR01000002.1 MAG TPA_asm: Candidatus Korarchaeota archaeon isolate HyVt-161
JAOZGN010000019.1 MAG: Candidatus Korarchaeota archaeon isolate INS_M22_B98
JALVGW010000419.1 MAG: Thermoproteota archaeon isolate A1_maxbin2_scaf2bin.274
JAOZFN010000006.1 MAG: Candidatus Korarchaeota archaeon isolate INS_M12_B43
JAOZFH010000026.1 MAG: Candidatus Korarchaeota archaeon isolate Aegir_M25_B147
JARZZN010000193.1+JARZZN010000222.1 MAG: Candidatus Korarchaeota archaeon isolate 17ROV19_HD25-17ROV19_HD25_MAG_00106
JAOZGE010000014.1 MAG: Candidatus Korarchaeota archaeon isolate INS_M21_B208
JAOZGP010000041.1 MAG: Candidatus Korarchaeota archaeon isolate INS_M34_B71
JAOZGM010000038.1 MAG: Candidatus Korarchaeota archaeon isolate INS_M22_B97
PNIB01000296.1 MAG: Candidatus Korarchaeota archaeon isolate ZAV-18
RCOS01000087.1 Candidatus Methanodesulfokores washburnensis strain MDKW
RXII01000074.1 MAG: Candidatus Methanodesulfokores washburnensis isolate NM4
QNVG01000086.1 MAG: Candidatus Korarchaeota archaeon isolate WYZ-LMO9
JAOZFG010000017.1 MAG: Candidatus Korarchaeota archaeon isolate Aegir_M25_B107
JALVGY010000442.1 MAG: Thermoproteota archaeon isolate A1_maxbin2_scaf2bin.286
JALXEK010000301.1 MAG: Thermoproteota archaeon isolate T2_metabat2_scaf2bin.391
MAIU01000107.1 MAG: Candidatus Korarchaeota archaeon NZ13-K Knz13_149
CALJEO010000156.1 MAG: uncultured archaeon isolate S1_Bin_METABAT__350_sub_1
JANXES010000117.1+JANXES010000171.1 MAG: Candidatus Korarchaeum sp. isolate SKYB86
JAVQRZ010000198.1 MAG: Candidatus Korarchaeum sp. isolate SKYBB_h_bin43
JAJHQZ010000001.1 MAG: Candidatus Korarchaeum sp. isolate Bin S22
DTEP01000052.1 MAG TPA_asm: Candidatus Korarchaeota archaeon isolate SpSt-718
DSGI01000089.1 MAG TPA_asm: Candidatus Korarchaeota archaeon isolate SpSt-15
RCOR01000014.1 Candidatus Korarchaeum cryptofilum strain WS Ga0080004_1113601
CP000968.1 Candidatus Korarchaeum cryptofilum OPF8
DTMV01000163.1 MAG TPA_asm: Candidatus Korarchaeota archaeon isolate SpSt-910
CALCBE010000124.1 MAG TPA_asm: Candidatus Korarchaeota archaeon isolate SRR6823441_bin.252_CONCOCT_v1.1_MAG
JAGGTZ010000019.1 MAG: Candidatus Korarchaeota archaeon isolate AUK061
JAOZFP010000020.1 MAG: Candidatus Korarchaeota archaeon isolate INS_M12_B58
JAOZFS010000007.1 MAG: Candidatus Korarchaeota archaeon isolate INS_M19_B79
JAOZFQ010000005.1 MAG: Candidatus Korarchaeota archaeon isolate INS_M13_B5
JAOZGR010000002.1 MAG: Candidatus Korarchaeota archaeon isolate INS_M9_B20
JAOZGK010000001.1 MAG: Candidatus Korarchaeota archaeon isolate INS_M22_B44
JAGHDG010000212.1 MAG: Candidatus Korarchaeota archaeon isolate AUK302
DQXR01000112.1 MAG TPA_asm: Candidatus Korarchaeota archaeon isolate HyVt-138
WALU01000059.1 MAG: Candidatus Korarchaeota archaeon isolate S016_48_esom
JALTGZ010000031.1 MAG: Thermoproteota archaeon isolate S016_metabat1_scaf2bin.017
JAOZFV010000038.1 MAG: Candidatus Korarchaeota archaeon isolate INS_M20_B204
JAVALW010000151.1 MAG: Candidatus Korarchaeota archaeon isolate 12ROV10_HD34E-12ROV10_HD34E_MAG_00031
JAVALV010000181.1+JAVALV010000059.1 MAG: Candidatus Korarchaeota archaeon isolate 12ROV10_HD34C-12ROV10_HD34C_MAG_00048
JAOZGJ010000017.1 MAG: Candidatus Korarchaeota archaeon isolate INS_M22_B125
JAOZFO010000022.1 MAG: Candidatus Korarchaeota archaeon isolate INS_M12_B44
JAOZFW010000080.1 MAG: Candidatus Korarchaeota archaeon isolate INS_M20_B56
JAOZGA010000154.1 MAG: Candidatus Korarchaeota archaeon isolate INS_M21_B149
JALUEI010000153.1 MAG: Thermoproteota archaeon isolate S143_metabat1_scaf2bin.045
WAOH01000220.1+WAOH0100021.1 MAG: Candidatus Korarchaeota archaeon isolate S143_77_esom
JALVIY010000189.1 MAG: Thermoproteota archaeon isolate A1_metabat1_scaf2bin.176
JALVNI010000008.1 MAG: Thermoproteota archaeon isolate A3_maxbin2_scaf2bin.223
JALSOH010000004.1 MAG: Thermoproteota archaeon isolate 356-284_metabat1_scaf2bin.099
JACXSD010000003.1 MAG: Candidatus Korarchaeota archaeon isolate 356-284-47
JACXSR010000024.1 MAG: Candidatus Korarchaeota archaeon isolate Hulk_76
JACXXK010000050.1 MAG: Candidatus Korarchaeota archaeon isolate PIR30_33
JAMOVA010000078.1 MAG: Thermoproteota archaeon isolate PIR-30_metabat1_scaf2bin.051
JARZZL010000570.1+JARZZL010000008.1 MAG: Candidatus Korarchaeota archaeon isolate 17ROV19_HD4-17ROV19_HD4_Bin_00097
JAVALX010000012.1 MAG: Candidatus Korarchaeota archaeon isolate 16ROV9_HD21-16ROV9_HD21_MAG_00046
JAOZFI010000043.1 MAG: Candidatus Korarchaeota archaeon isolate Aegir_M25_B233
JARZZM010000008.1 MAG: Candidatus Korarchaeota archaeon isolate 17ROV19_HD25-17ROV19_HD25_MAG_00082
JALVZR010000016.1 MAG: Thermoproteota archaeon isolate T10_metabat1_scaf2bin.052
JALVPC010000003.1 MAG: Thermoproteota archaeon isolate A3_metabat1_scaf2bin.054
JALWYH010000019.1 MAG: Thermoproteota archaeon isolate T2_maxbin2_scaf2bin.524
JALVFV010000076.1 MAG: Thermoproteota archaeon isolate A1_maxbin2_scaf2bin.056

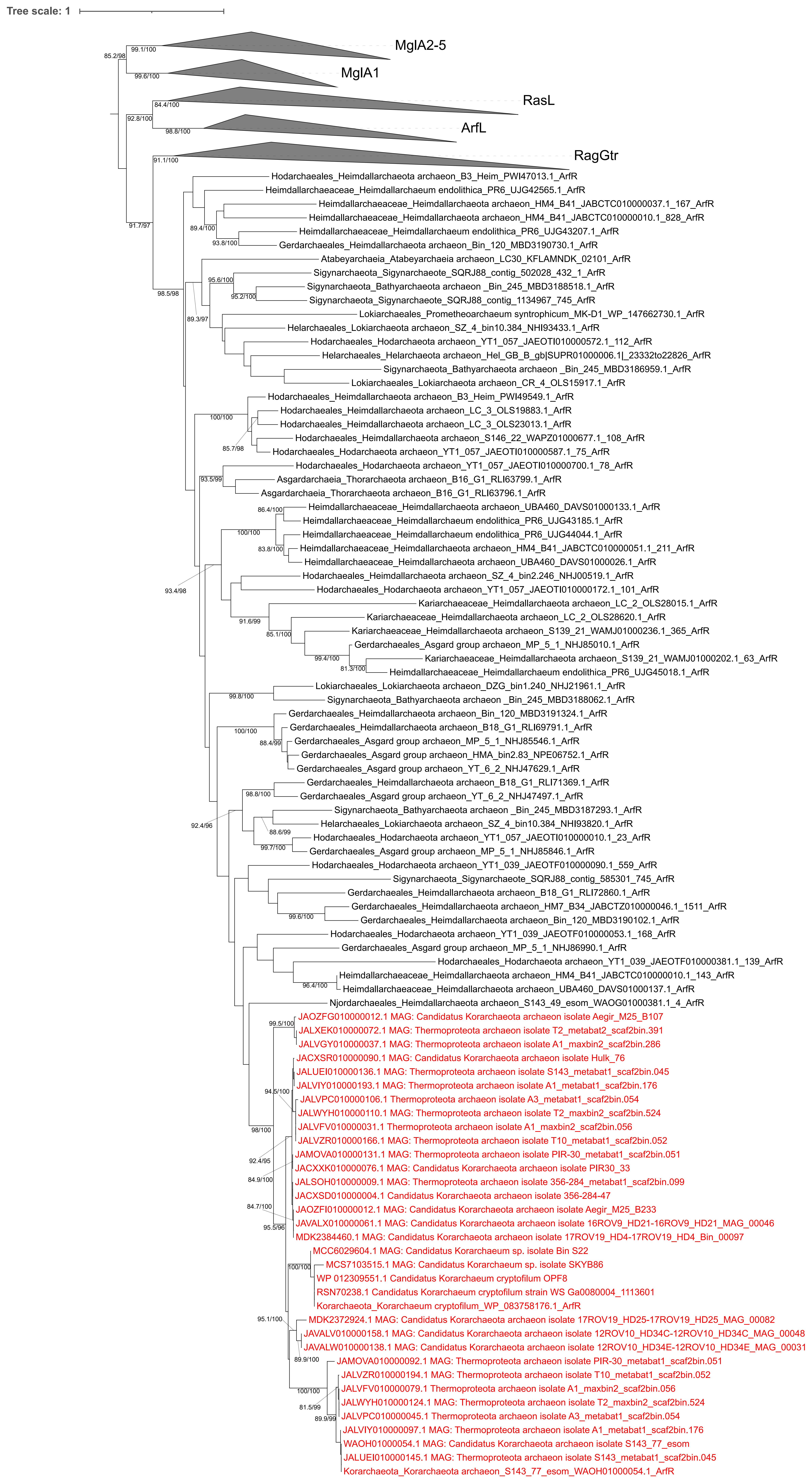
Tree scale: 1
MglA2-5
MglA1
RasL
ArfL
RagGtr
Hodarchaeales_Heimdallarchaeota archaeon_B3_Heim_PWI47013.1_ArfR
Heimdallarchaeaceae_Heimdallarchaeum endolithica_PR6_UJG42565.1_ArfR
Heimdallarchaeaceae_Heimdallarchaeota archaeon_HM4_B41_JABCTC010000037.1_167_ArfR
Heimdallarchaeaceae_Heimdallarchaeota archaeon_HM4_B41_JABCTC010000010.1_828_ArfR
Heimdallarchaeaceae_Heimdallarchaeum endolithica_PR6_UJG43207.1_ArfR
Gerdarchaeales_Heimdallarchaeota archaeon_Bin_120_MBD3190730.1_ArfR
Atabeyarchaeia_Atabeyarchaeia archaeon_LC30_KFLAMNDK_02101_ArfR
Sigynarchaeota_Sigynarchaeote_SQRJ88_contig_502028_432_1_ArfR
Sigynarchaeota_Bathyarchaeota archaeon _Bin_245_MBD3188518.1_ArfR
Sigynarchaeota_Sigynarchaeote_SQRJ88_contig_1134967_745_ArfR
Lokiarchaeales_Prometheoarchaeum syntrophicum_MK-D1_WP_147662730.1_ArfR
Helarchaeales_Lokiarchaeota archaeon_SZ_4_bin10.384_NHI93433.1_ArfR
Hodarchaeales_Hodarchaeota archaeon_YT1_057_JAEOTI010000572.1_112_ArfR
Helarchaeales_Helarchaeota archaeon_Hel_GB_B_gb|SUPR01000006.1|_23332to22826_ArfR
Sigynarchaeota_Bathyarchaeota archaeon _Bin_245_MBD3186959.1_ArfR
Lokiarchaeales_Lokiarchaeota archaeon_CR_4_OLS15917.1_ArfR
Hodarchaeales_Heimdallarchaeota archaeon_B3_Heim_PWI49549.1_ArfR
Hodarchaeales_Heimdallarchaeota archaeon_LC_3_OLS19883.1_ArfR
Hodarchaeales_Heimdallarchaeota archaeon_LC_3_OLS23013.1_ArfR
Hodarchaeales_Heimdallarchaeota archaeon_S146_22_WAPZ01000677.1_108_ArfR
Hodarchaeales_Hodarchaeota archaeon_YT1_057_JAEOTI010000587.1_75_ArfR
Hodarchaeales_Hodarchaeota archaeon_YT1_057_JAEOTI010000700.1_78_ArfR
Asgardarchaeia_Thorarchaeota archaeon_B16_G1_RLI63799.1_ArfR
Asgardarchaeia_Thorarchaeota archaeon_B16_G1_RLI63796.1_ArfR
Heimdallarchaeaceae_Heimdallarchaeota archaeon_UBA460_DAVS01000133.1_ArfR
Heimdallarchaeaceae_Heimdallarchaeum endolithica_PR6_UJG43185.1_ArfR
Heimdallarchaeaceae_Heimdallarchaeum endolithica_PR6_UJG44044.1_ArfR
Heimdallarchaeaceae_Heimdallarchaeota archaeon_HM4_B41_JABCTC010000051.1_211_ArfR
Heimdallarchaeaceae_Heimdallarchaeota archaeon_UBA460_DAVS01000026.1_ArfR
Hodarchaeales_Heimdallarchaeota archaeon_SZ_4_bin2.246_NHJ00519.1_ArfR
Hodarchaeales_Hodarchaeota archaeon_YT1_057_JAEOTI010000172.1_101_ArfR
Kariarchaeaceae_Heimdallarchaeota archaeon_LC_2_OLS28015.1_ArfR
Kariarchaeaceae_Heimdallarchaeota archaeon_LC_2_OLS28620.1_ArfR
Kariarchaeaceae_Heimdallarchaeota archaeon_S139_21_WAMJ01000236.1_365_ArfR
Gerdarchaeales_Asgard group archaeon_MP_5_1_NHJ85010.1_ArfR
Kariarchaeaceae_Heimdallarchaeota archaeon_S139_21_WAMJ01000202.1_63_ArfR
Heimdallarchaeaceae_Heimdallarchaeum endolithica_PR6_UJG45018.1_ArfR
Lokiarchaeales_Lokiarchaeota archaeon_DZG_bin1.240_NHJ21961.1_ArfR
Sigynarchaeota_Bathyarchaeota archaeon _Bin_245_MBD3188062.1_ArfR
Gerdarchaeales_Heimdallarchaeota archaeon_Bin_120_MBD3191324.1_ArfR
Gerdarchaeales_Heimdallarchaeota archaeon_B18_G1_RLI69791.1_ArfR
Gerdarchaeales_Asgard group archaeon_MP_5_1_NHJ85546.1_ArfR
Gerdarchaeales_Asgard group archaeon_HMA_bin2.83_NPE06752.1_ArfR
Gerdarchaeales_Asgard group archaeon_YT_6_2_NHJ47629.1_ArfR
Gerdarchaeales_Heimdallarchaeota archaeon_B18_G1_RLI71369.1_ArfR
Gerdarchaeales_Asgard group archaeon_YT_6_2_NHJ47497.1_ArfR
Sigynarchaeota_Bathyarchaeota archaeon _Bin_245_MBD3187293.1_ArfR
Helarchaeales_Lokiarchaeota archaeon_SZ_4_bin10.384_NHI93820.1_ArfR
Hodarchaeales_Hodarchaeota archaeon_YT1_057_JAEOTI010000010.1_23_ArfR
Gerdarchaeales_Asgard group archaeon_MP_5_1_NHJ85846.1_ArfR
Hodarchaeales_Hodarchaeota archaeon_YT1_039_JAEOTF010000090.1_559_ArfR
Sigynarchaeota_Sigynarchaeote_SQRJ88_contig_585301_745_ArfR
Gerdarchaeales_Heimdallarchaeota archaeon_B18_G1_RLI72860.1_ArfR
Gerdarchaeales_Heimdallarchaeota archaeon_HM7_B34_JABCTZ010000046.1_1511_ArfR
Gerdarchaeales_Heimdallarchaeota archaeon_Bin_120_MBD3190102.1_ArfR
Hodarchaeales_Hodarchaeota archaeon_YT1_039_JAEOTF010000053.1_168_ArfR
Gerdarchaeales_Asgard group archaeon_MP_5_1_NHJ86990.1_ArfR
Hodarchaeales_Hodarchaeota archaeon_YT1_039_JAEOTF010000381.1_139_ArfR
Heimdallarchaeaceae_Heimdallarchaeota archaeon_HM4_B41_JABCTC010000010.1_143_ArfR
Heimdallarchaeaceae_Heimdallarchaeota archaeon_UBA460_DAVS01000137.1_ArfR
Njordarchaeales_Heimdallarchaeota archaeon_S143_49_esom_WAOG01000381.1_4_ArfR
JAOZFG010000012.1 MAG: Candidatus Korarchaeota archaeon isolate Aegir_M25_B107
JALXEK010000072.1 MAG: Thermoproteota archaeon isolate T2_metabat2_scaf2bin.391
JALVGY010000037.1 MAG: Thermoproteota archaeon isolate A1_maxbin2_scaf2bin.286
JACXSR010000090.1 MAG: Candidatus Korarchaeota archaeon isolate Hulk_76
JALUEI010000136.1 MAG: Thermoproteota archaeon isolate S143_metabat1_scaf2bin.045
JALVIY010000193.1 MAG: Thermoproteota archaeon isolate A1_metabat1_scaf2bin.176
JALVPC010000106.1 Thermoproteota archaeon isolate A3_metabat1_scaf2bin.054
JALWYH010000110.1 MAG: Thermoproteota archaeon isolate T2_maxbin2_scaf2bin.524
JALVFV010000031.1 Thermoproteota archaeon isolate A1_maxbin2_scaf2bin.056
JALVZR010000166.1 MAG: Thermoproteota archaeon isolate T10_metabat1_scaf2bin.052
JAMOVA010000131.1 MAG: Thermoproteota archaeon isolate PIR-30_metabat1_scaf2bin.051
JACXXK010000076.1 MAG: Candidatus Korarchaeota archaeon isolate PIR30_33
JALSOH010000009.1 MAG: Thermoproteota archaeon isolate 356-284_metabat1_scaf2bin.099
JACXSD010000004.1 Candidatus Korarchaeota archaeon isolate 356-284-47
JAOZFI010000012.1 MAG: Candidatus Korarchaeota archaeon isolate Aegir_M25_B233
JAVALX010000061.1 MAG: Candidatus Korarchaeota archaeon isolate 16ROV9_HD21-16ROV9_HD21_MAG_00046
MDK2384460.1 MAG: Candidatus Korarchaeota archaeon isolate 17ROV19_HD4-17ROV19_HD4_Bin_00097
MCC6029604.1 MAG: Candidatus Korarchaeum sp. isolate Bin S22
MCS7103515.1 MAG: Candidatus Korarchaeum sp. isolate SKYB86
WP 012309551.1 Candidatus Korarchaeum cryptofilum OPF8
RSN70238.1 Candidatus Korarchaeum cryptofilum strain WS Ga0080004_1113601
Korarchaeota_Korarchaeum cryptofilum_WP_083758176.1_ArfR
MDK2372924.1 MAG: Candidatus Korarchaeota archaeon isolate 17ROV19_HD25-17ROV19_HD25_MAG_00082
JAVALV010000158.1 MAG: Candidatus Korarchaeota archaeon isolate 12ROV10_HD34C-12ROV10_HD34C_MAG_00048
JAVALW010000138.1 MAG: Candidatus Korarchaeota archaeon isolate 12ROV10_HD34E-12ROV10_HD34E_MAG_00031
JAMOVA010000092.1 MAG: Thermoproteota archaeon isolate PIR-30_metabat1_scaf2bin.051
JALVZR010000194.1 MAG: Thermoproteota archaeon isolate T10_metabat1_scaf2bin.052
JALVFV010000079.1 Thermoproteota archaeon isolate A1_maxbin2_scaf2bin.056
JALWYH010000124.1 MAG: Thermoproteota archaeon isolate T2_maxbin2_scaf2bin.524
JALVPC010000045.1 Thermoproteota archaeon isolate A3_metabat1_scaf2bin.054
JALVIY010000097.1 MAG: Thermoproteota archaeon isolate A1_metabat1_scaf2bin.176
WAOH01000054.1 MAG: Candidatus Korarchaeota archaeon isolate S143_77_esom
JALUEI010000145.1 MAG: Thermoproteota archaeon isolate S143_metabat1_scaf2bin.045
Korarchaeota_Korarchaeota archaeon_S143_77_esom_WAOH01000054.1_ArfR
